# Homologous ABA-independent kinase tracks coalesced into osmotic stress circuits during plant terrestrialization

**DOI:** 10.64898/2026.05.21.726866

**Authors:** Jaccoline M.S. Zegers, Gabriel X. Garcia Ramirez, Anne Harzen, Sara C. Stolze, Mohamed Salem, Anne Ziplys, Isabel Mora-Ramírez, Michal Shpilman, Assaf Mosquna, Kerstin Schmitt, Craig A. Dorsch, Cäcilia F. Kunz, Stefanie König, Leo Hofmann, Gerhard H. Braus, Ivo Feussner, Oliver Valerius, Juan C. Moreno, Salim Al-Babili, Jos H.M. Schippers, Hirofumi Nakagami, Jan de Vries

## Abstract

Land plants possess a unique system for responding to environmental stressors. How this system evolved during plant terrestrialization remains one of the major questions in plant evolutionary biology. To retrace this process, it is essential to study both land plants and their closest algal relatives, the zygnematophytes. Using the single-celled zygnematophyte *Mesotaenium*, we integrated physiological stress experiments with phosphoproteomics, genome-wide transcription factor binding analyses, and protein–protein interaction studies to investigate the architecture of a key stress response pathway: the signaling cascade homologous to the plant abscisic acid (ABA)-mediated pathway. Our results highlight the roles of histidine kinases (HKs) and calcium-dependent protein kinases (CDPKs) in osmotic stress signaling. Focusing on SnRK2 and ABF, key components at the downstream end of the canonical ABA signaling pathway, we provide evidence that ABF plays a central role in osmotic stress responses even in the absence of ABA. Together, our data reveal the coordinated action of parallel functional modules that were likely integrated into the ABA response cascade during plant terrestrialization.

## INTRODUCTION

Stress abounds on land. In land plants, a powerful genetic regulatory network integrates the perception of the diverse challenging cues and mounts a suite of protective responses^1,2^. In both land plants and their algal relatives include the synthesis of osmoprotective proteins and metabolites – such as Late Embryogenesis Abundant (LEA) proteins^3^, proline and more chemodiverse metabolites^4–9^ – as well as the regulation of water and ion transport systems^10^.

Central to this acclimation response is the well-characterized abscisic acid (ABA) signaling cascade^11,12^ that is found across land plants^13–19^. Under drought, salinity, or cold stress – each of which involves a (pre-)hyperosmotic component – ABA levels increase either through de novo synthesis or redistribution. ABA then binds to PYR/PYL/RCAR receptors enabling them to inhibit type 2C protein phosphatases (PP2C, particularly clade A)^20–24^. Under non-stress conditions, these phosphatases inhibit SnRK2 kinases. However, upon ABA perception and PP2CA inhibition, SnRK2s are no longer suppressed and can undergo lasting (auto-)phosphorylation in their activation loop, becoming catalytically active. Once activated, SnRK2 kinases phosphorylate a range of downstream targets. One key substrate is the transcription factor ABF (ABA-responsive element Binding Factor)^25^, which in turn regulates the expression of stress-responsive genes, including LEA^26,27^. SnRK2s also phosphorylate ion channels such as Slow Anion Channel 1 (SLAC1), contributing to stomatal closure and reducing water loss^28–30^. More recently, RAF-like kinases have emerged as additional upstream activators of SnRK2^31,32^, for some even required for phosphorylation of their activation loops, highlighting a complex regulatory network. In the moss *Physcomitrium patens*, for example, RAF kinases play a pivotal role in ABA signaling, positioning SnRK2 as a central regulatory node.

The evolutionary origins of this pathway have gained increasing attention, especially following the discovery of homologs to PYL proteins in certain orders of Zygnematophyceae^33–35^ – the algal sister group to embryophytes^36,37^ (land plants). Interestingly, these algal *PYL* homologs co-express with other genes of the stress module^38^, their transcripts change under stress^4,39^, and these zygnematophyte PYL proteins can inhibit PP2C phosphatases in an ABA-independent manner^39^. This raises important questions about this signaling module. How does this pathway operate in the absence of ABA?^40^ While no other algal group has PYL homologs, other key components such as SnRK2, PP2C, ABF, and RAF are present and widely conserved across streptophytes. How did these assemble into the signaling module of ABA perception as we know it in today’s embryophytes?

To reconstruct the early evolutionary history of this signaling module, we investigated the stress signaling responses of *Mesotaenium endlicherianum*, a unicellular zygnematophycean alga, under hyperosmotic and cold stress conditions. Our particular focus was on the roles of SnRK2 and ABF in these responses. By studying this alga – an extant relative of early land plant ancestors – we aim to gain insight into how key elements of the ABA signaling cascade functioned over 500 million years ago, prior to the colonization of land.

## RESULTS AND DISCUSSION

### A phylogenetic framework for the osmotic stress response module

In land plants, the core ABA module is formed out of PYL, PP2C, SnRK2 and downstream targets. SnRK2 sequences are highly conserved across streptophyte algae, with most algal species typically harboring a single copy. Notably, every streptophyte alga in our dataset possessed this gene, suggesting that SnRK2 plays an essential and conserved role in these organisms. This pattern changes markedly in Tracheophytes – and possibly in some Bryophytes – where multiple SnRK2 subgroups are present, enabeling subfunctionalization and neofunctionalization. Determining which functions are ancestral and which are novel remains unresolved. Simple comparisons between subgroups are insufficient to draw definitive conclusions about their evolutionary trajectories. Overall, we resolved relationships of SnRK2s among more recently diverged lineages with high support while the backbone support for the deep genetic structure was low – except for the consistent clustering of Zygnematophyceae (Fig. 1a). To enhance the reliability of the deep branches in our phylogenetic tree, we increased representation from streptophyte algae and reduced the number of embryophyte sequences rampant with neofunctionalization and subfunctionalization. For instance, only subgroup III is specifically responsible for ABA-regulated stomatal closure. Simultaneously, substantial functional redundancy exists among the subgroups, as knocking out 9 of the 10 SnRK2 genes in *Arabidopsis* does not result in a severe phenotype, provided SnRK2.2 remains active^41^.

**Figure 1.**
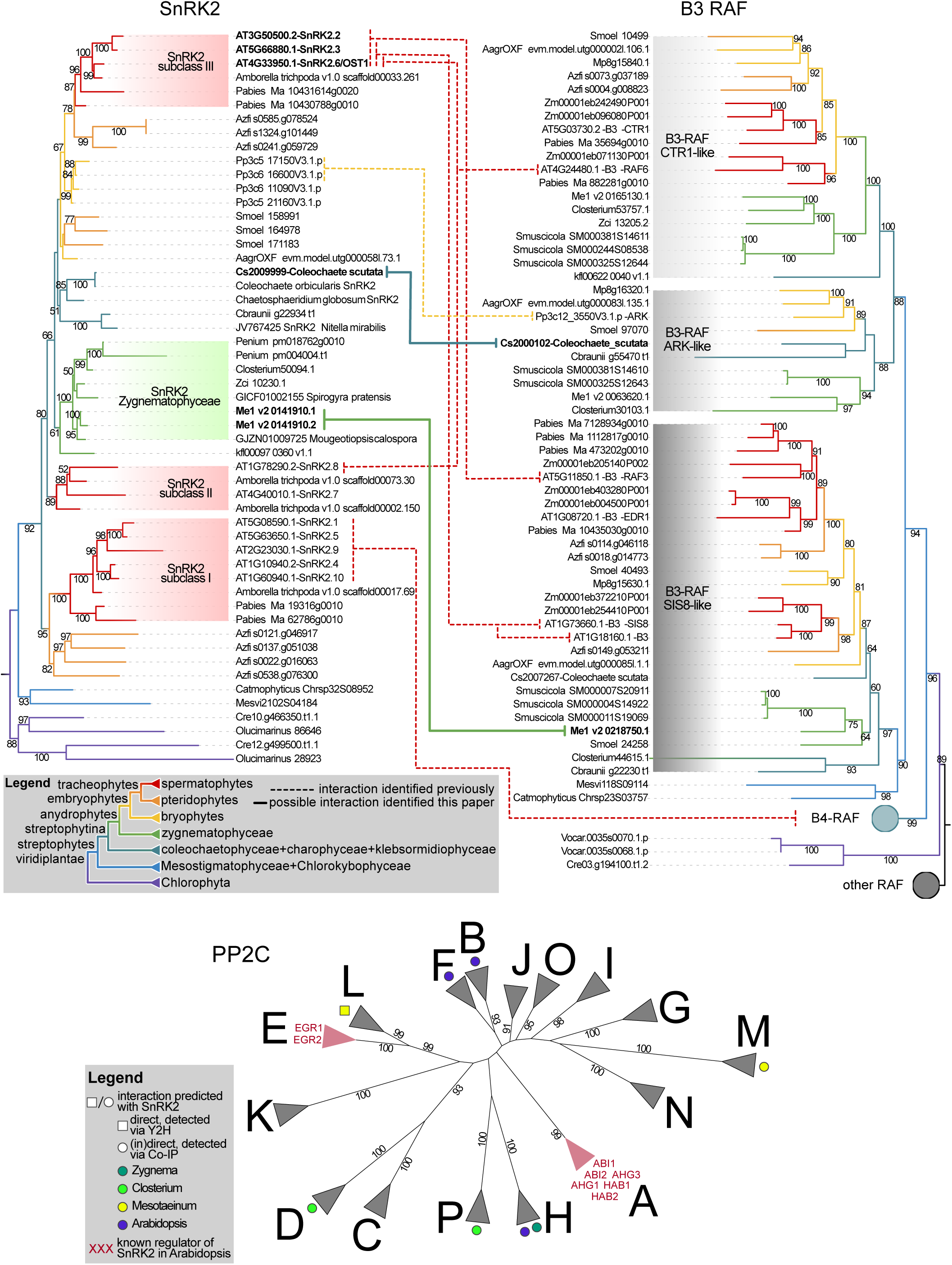
Phylogenetic analysis of SnRK2, B3-RAF, and PP2C proteins across streptophytes. Phylogenetic trees were constructed for SnRK2, B3-RAF, and PP2C protein families identified across representative streptophyte species. For SnRK2, previously defined subgroups from *Arabidopsis* thaliana are indicated. B3-RAF proteins were assigned to newly defined subgroups based on this analysis. Dashed lines represent previously reported protein-protein interactions between SnRK2 and B3-RAF proteins, while solid lines denote potential novel interactions identified in this study. PP2C proteins were grouped according to clades A-L, as defined by Bhaskara et al.^42^; additional clades M-P were established here to categorize sequences that did not align with existing groupings.

Previous phylogenetic studies have classified RAF kinases into distinct groups; we focused on the B3-RAF clade. B3-RAF kinases cluster most closely with B4-RAF kinases. Notably, B3/B4-RAF kinases from Chlorophyta fall outside both the B3 and B4 clades, suggesting that the divergence between B3 and B4 kinases likely occurred in the last common ancestor (LCA) of the streptophytes. Within the B3-RAF clade, we identified three subgroups that reflect the diversity of streptophytic lineages: the CTR1-like, ARK-like, and SIS8-like subgroups – named after well-characterized *Physcomitrium* and *Arabidopsis* proteins. All streptophytes in our dataset possess at least one B3-RAF kinase, though not necessarily a representative for each subgroup.

We recovered all established^42^ subgroups of PP2Cs, clades A to L. All known *Arabidopsis* PP2Cs clustered within these expected clades, with the exception of the pseudophosphatase DOG18, which grouped with clade A. For previously unclassified *Arabidopsis* PP2Cs, we established three novel clades – M, N, and O. Additionally, we identified one clade lacking any *Arabidopsis* representatives, which we designated as clade P. Based on bootstrap support, clade F should be subdivided into three smaller clades. Similarly, clades E and K could also be further divided, as multiple copies appear to have existed already in the LCA of the Viridiplantae. However, such subdivision would not be necessary when focusing solely on *Arabidopsis*.

### Quantitive phosphoproteome profiling identifies conserved osmotic stress proteins

Within the canonical ABA signaling phosphorelay, SnRK2 is activated by phosphorylation, and once activated phosphorylates ABF transcription factors^43–46^. We thus investigated the phosphoproteome (S/T/Y) dynamics of *Mesotaenium* following hyperosmotic treatments for 1h and 3h (Fig. 2a-b). As expected, a substantial portion of the phosphoproteome is derived from signaling proteins, with 19% of the identified phosphopeptides originating from protein kinases and transcriptional-associated proteins^47^ (TAPs). Interestingly, 22% of the phosphoproteome corresponds to proteins of unknown function, highlighting the extent of unexplored regulatory mechanisms. We quantified over 6,741 and 6,474 phosphopeptides per time point; 683 and 474 phosphopeptides were classified as changing in their abundance among the treatments based on a reliability-adjusted F-score (RAF-score; a penalized ANOVA F-statistic accounting for the extent and distribution of imputed values). A cutoff of RAF-score ≥ 7 was chosen, which would correspond to p ≲ 0.001 for an individual (no multiple-testing correction), true F-statistic. Most of the variance in the data could be attributed to the treatment conditions (salt, mannitol, cold, and control), rather than the time points (Fig. 2b and Supplementary Fig. 1b). Notably, the two hyperosmotic treatments (salt and mannitol) clustered most closely together along the first two principal components and showed the strongest correlation among treatments.

**Figure 2.**
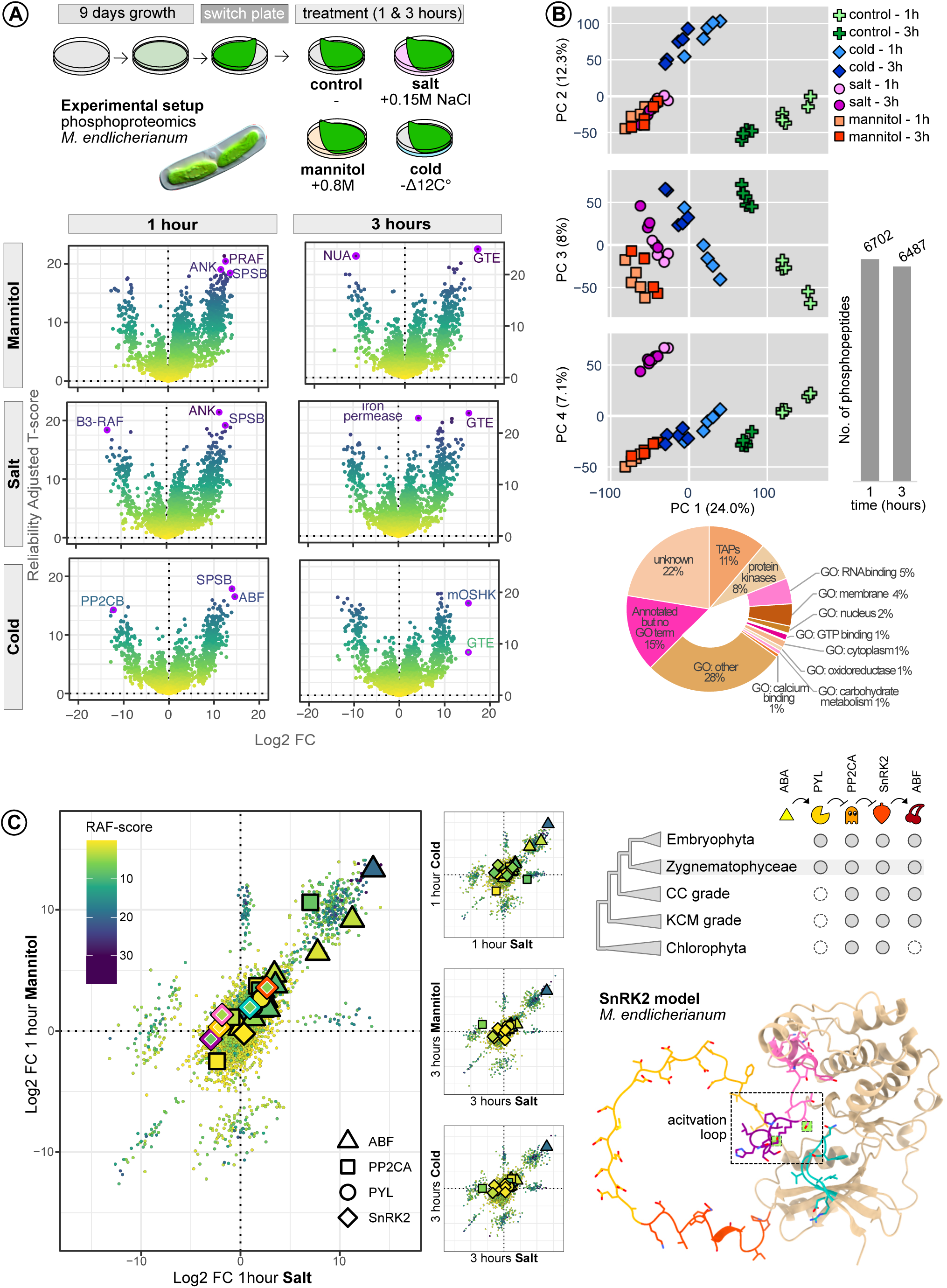
Phosphoproteomics of *Mesotaenium endlicherianum* under hyperosmotic and cold stress. A) Overview of the experimental design and volcano plots displaying all identified phosphopeptides. Protein abbreviations; ANK = Ankyrin repeat protein; B3-RAF = RAF protein kinase class B3; PRAF = Photosynthesis-related RAF-like protein kinase; SPSB = Sucrose-Phosphate Synthase B; PP2CB = protein phosphatase 2C class B; ABF = ABA-responsive element binding factor; NUA = Nuclear-pore anchor; GTE = Global Transcription Factor of Group E. B) Exploratory data analysis, including principal component analysis (PCA) across the first four principal components, and the distribution of functional protein annotations among the identified peptides. C) Identified phosphopeptides associated with the canonical ABA signaling cascade. A structural model of SnRK2 is shown on the right, with each quantified phosphopeptide highlighted in a distinct color. The serine residues homologues to SER171 and SER175 in *At*SnRK2.6 are highlighted in green. FC = Fold Change. RAF-score = Reliability adjusted F-score, which is based on the ANOVA F-statistic adjusted for number of missing values, see method section for more details.

For the comparison across treatments (Fig. 2a,c, 3b; see also the Shiny App), two key behaviors of the phosphoproteome must be considered. First, phosphopeptides can be consistently detected across conditions while varying in abundance. Second, peptides may transition between unphosphorylated and phosphorylated states (and vice versa) between treatments, resulting in apparent large fold changes due to missing values in one state. As four conditions are compared simultaneously, combinations of these two behaviors for one peptide can also occur. Among the differentially abundant phosphopeptides, many are associated with well-known signaling proteins such as kinases, transcription factors, and RNA-modifying enzymes. Additionally, we identified phosphorylation changes in proteins known to be involved in hyperosmotic stress responses, including sucrose-phosphate synthase (involved in osmolyte biosynthesis), ion channels (regulating ion homeostasis), and vacuolar sorting proteins.

**Figure 3.**
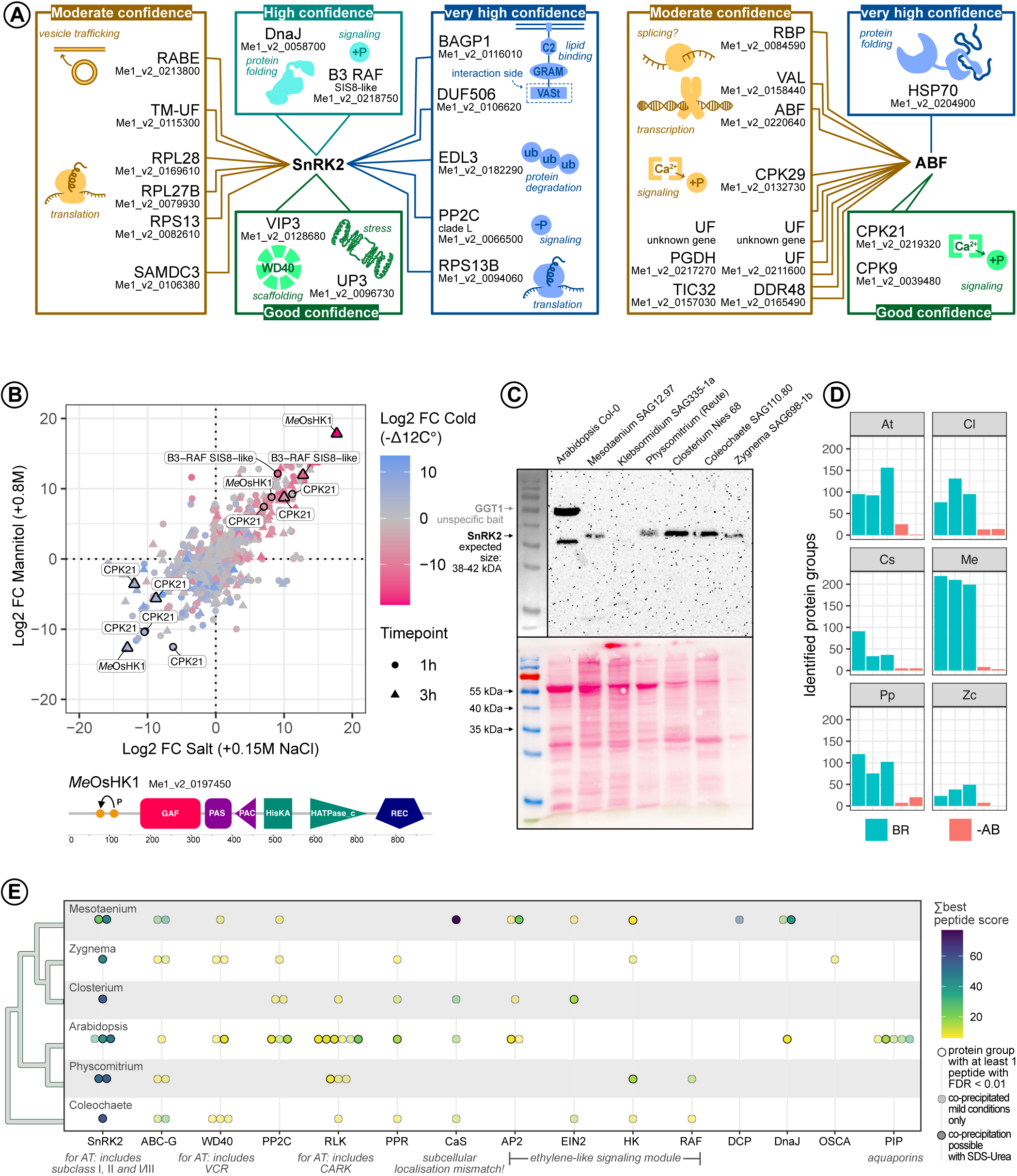
Identification of SnRK2 and ABF interactors in *Mesotaenium endlicherianum*. A) Yeast two-hybrid screening results using SnRK2 (left) and ABF (right) as bait proteins. Confidence levels are based on both the number and probability of detected interactions. B) Log2 fold changes of phosphopeptides corresponding to *Mesotaenium* kinases. Highlighted are a potential RAF kinase interactor of SnRK2 and a possible CPK21 interactor of ABF, along with *Me*OsHK1 (*Mesotaenium* Osmotic Stress Histidine Kinase 1), the most significantly phosphoregulated kinase. Domain architecture and dominant phosphosites of *Me*OsHK1 are shown. C-E) Co-IP experiments across streptophyte species using SnRK2 as bait. C) Western blot of total protein extracts (top) and blot probed with SnRK2-specific antibody (PHY0715A), ponceau stain below. D) Number of protein groups identified per sample. BR = biological replicate; −AB = no antibody control. E) Protein groups identified in more than three species and absent in all −AB controls, previously reported as key components of hyperosmotic stress or SnRK2 signaling pathways.

We next scrutinized phosphopeptides corresponding to each component of the core ABA signaling cascade (Fig. 2c). If this pathway functions in *Mesotaenium* similarly to how it does in *Arabidopsis*, we would expect phospho-regulation of both *Me*SnRK2 and *Me*ABF. Of the 11 *Me*ABF-associated phosphopeptides detected, all but one exhibited increased abundance upon salt, mannitol, and cold treatment (at sites distinct from those identified in *Arabidopsis*^44^), although responses differed markedly in magnitude and statistical support. *Me*ABF’s primary phosphopeptide was the highest-scoring TAP phosphopeptide across all treatments at 1h. After 3h, *Me*ABF phosphorylation remained elevated under both hyperosmotic and cold treatments, but was surpassed by a phosphopeptide from another transcription factor, Global Transcription Factor Group E4 (*Me*GTE4) in all three conditions, which is likely involved in cell division and chromatin remodeling^48^. For SnRK2, we identified at least five phosphorylatable sites (Fig. 2c). If the kinase were activated similarly to *Arabidopsis*, phosphorylation within the activation loop would be expected. In *Arabidopsis*, the activation loop of *At*SnRK2.6 contains two phosphorylation sites, SER171 and SER175, both of which contribute to kinase activation^49^. However, the first corresponding phosphopeptide, *Me*SSLLHSQPK+p (↔ *At*SnRK2.6 SSVLHpSQPK; SER171), decreased in abundance upon salt stress, contrary to expectations, and remained unchanged under the other treatments. The second phosphopeptide from the activation loop, *Me*STVGTPAYIAPEVLSK+p (↔ *At*SnRK2.6 pSTVGTPAYIAPEVLLK; SER175), exhibited minor variation in abundance among treatments, although these changes did not exceed the applied RAF-score threshold. Additionally, we detected phosphorylation of PYL and PP2CA components. Among these, only one site in PP2CA showed a significant change in phosphorylation upon treatments.

Following these observations, it is plausible to hypothesize that *Me*ABF does get activated upon osmotic stress in *Mesotaenium*. For *Me*SnRK2, the situation remains unclear. We did not observe hyperphosphorylation of the activation loop during osmotic stress relative to the control treatment, pointing into the direction that SnRK2 was not activated during any of the treatments. It is also possible that *Me*SnRK2 activation occurred too transiently to capture experimentally, or that the kinase was already partially or fully active before the treatment began.

### Genome-wide SnRKs and ABF interactions partners

To further elucidate the roles of ABF and SnRK2 in *Mesotaenium*, we performed a genome-wide yeast two-hybrid (Y2H) screen against a library of all expressed protein-coding genes (Fig. 3a). No interaction was detected between ABF and SnRK2, in either direction, opposite of what would be expected if ABF were a substrate of SnRK2. Consistent with data on land plants^50^, we identified calcium-dependent protein kinases (CDPKs) as potential ABF regulators. Osmotic stress can trigger a rapid influx of calcium via stretch-activated ion channels such as OSCAs^51^ (Hyperosmolarity-Gated Ca²⁺-Permeable Channels) and other calcium channels activated through cell wall-associated receptors. Such calcium influxes lead to activation of CDPKs, which in turn could phosphorylate downstream transcription factors like ABF as it occurs in land plants^52,53^. One of the identified CDPKs (*Me*CPK21) showed dynamic phosphor-regulation, with several phosphosites showing either strongly phosphorylation or dephosphorylation in response to osmotic stress (Fig. 3b). CDPKs are known to undergo autophosphorylation upon calcium binding, which can modulate their activity, stability, or interactions with other proteins. However, it is important to note that autophosphorylation is not strictly required for activation; calcium-induced conformational changes alone can suffice to activate CDPKs.

In addition to CDPKs, HSP70 emerged as a highly likely interactor of ABF in the yeast two-hybrid screen. While the biological relevance of this interaction remains speculative, it is possible that HSP70 supports ABF folding or stabilization, as is common for molecular chaperones. We also identified several moderate-confidence interactors, one notable hit being ABF itself, aligning with this leucine zipper forming homodimers^26^.

The Y2H screen yielded more, and stronger, candidate interactors for SnRK2 than for ABF. Among these were the expected interactors: PP2C and B3-RAF^54^. PP2Cs negatively regulate SnRK2 by dephosphorylation. In particular, clade A PP2Cs are key repressors in the canonical ABA signaling pathway, while clade E PP2Cs have been implicated in SnRK2 regulation under cold stress^55,56^. However, the PP2C identified here belongs to clade L (Fig. 1) – not previously associated with SnRK2 regulation. Clade L is phylogenetically the closest to clade E, suggesting it may serve a similar function in *Mesotaenium*. SnRK2 proteins have over 80% sequence similarity between Zygnematophyceae and embryophytes (and >50% between chlorophytes and streptophytes). Thus, to explore conserved SnRK2 interactors across streptophytes, we performed co-immunoprecipitation (Co-IP) using the same SnRK2 antibody on six phragmoplatophytes covering more than 730 million years of evolutionary divergence^57^ and *Klebsormidium* as outgroup. We detected a specific SnRK2 band in all but *Klebsormidium* in initial Western blot analysis (Fig. 3c) and thus excluded *Klebsormidium* from further Co-IPs. In all Co-IPs we successfully pulled down SnRK2. Our exploratory Co-IP revealed potential PP2C interactors in four out of six species – almost all from different clades (Fig. 1). These findings suggest that SnRK2 regulation by PP2Cs may be evolutionarily widespread but more flexible than previously thought. Furthermore, we recovered several known interactors (Fig. 3d) – either previously reported or identified in our Y2H screen. Notably, however, we did not detect ABF or PP2CA in any of the samples.

B3-RAF kinases have recently gained increased attention as positive regulators of SnRK2 signaling in both *Physcomitrium* and *Arabidopsis*^13,58^. In *Arabidopsis*, this role likely involves multiple B3-RAF family members. B3-RAFs belong to a subclass of RAF kinases, which themselves are a subgroup within the MAPKKK (mitogen-activated protein kinase kinase kinase) family. Notably, well-known B3-RAFs such as Constitutive Triple Response 1 (CTR1), a key player in ethylene signaling^59^, and Enhanced Disease Resistance 1 (EDR1), involved in pathogen response^60^, are not directly upstream of SnRK2. However, several closely related B3-RAFs are implicated in SnRK2 regulation (Fig. 1). In *Physcomitrium*, there is only one known B3-RAF, abiotic stress-responsive Raf-like kinase (ARK), which positively regulates SnRK2 within the ABA pathway^61^. ARK is itself regulated by an Ethylene Response Histidine Kinase (ETR-HK), and disruption of this ETR-HK abolishes ABA signaling entirely. Remarkably, the ETR-HK in *Physcomitrium* can be functionally replaced by homologs from other bryophytes and the streptophyte algae *Klebsormidium*, but not by the *Arabidopsis* ETR-HK, indicating divergence in upstream regulatory mechanisms. In *Arabidopsis*, the upstream activators of B3-RAFs remain unknown. However, these kinases are rapidly activated – within minutes – following osmotic stress^62^, suggesting their role in swift signaling events.

In *Mesotaenium*, we identified a B3-RAF belonging to the SIS8 clade (Fig. 1) phosphorylated upon osmotic stress (Fig. 3b), indicating functional activation. Our cross-species Co-IP screen further identified potential RAF kinase interactors of SnRK2: a B1-clade RAF kinase in *Physcomitrium* and a B3-ARK-like kinase in *Coleochaete* (Figs. 1, 3e). These findings suggest that the regulatory interaction between SnRK2 and RAF kinases predates the emergence of land plants, pointing to an ancient and conserved osmotic stress signaling loop.

Within the *Mesotaenium* kinome (Fig. 3b), the phosphopeptides showing the most pronounced changes mapped to a histidine kinase (HK). While the top BLASTp hit for *Arabidopsis* was ETHYLENE RESPONSE 1 (ETR1), the domain architecture of the *Mesotaenium* kinase diverges by lacking the characteristic transmembrane domains. Instead, phosphorylation events were concentrated in the N-terminal sensory region – an unusual region that appears to be unique to *Mesotaenium*, pointing to a putative unknown zygnematophycean signaling system. Phylogenetic analysis further revealed that this HK belongs to a clade of HKs that has been lost altogether in *Arabidopsis* and other land plants (Supplementary Fig. 2). This HK was not an isolated case: a broad array of HKs showed strong phosphorylation responses to hyperosmotic stress, suggesting that HK-mediated signaling plays a substantial role in osmotic stress adaptation in *Mesotaenium*.

In the predicted *Mesotaenium* SnRK2 interactome, several ribosomal proteins appeared as putative SnRK2 interactors. This raises the intriguing possibility that SnRK2 might also directly influence translation in *Mesotaenium* next to transcription. Additionally, we identified a homolog of BAG-Associated GRAM Protein 1 (BAGP1)^63^, a moderately characterized protein in *Arabidopsis*. Our results predict that *Mesotaenium* BAGP1 interacts with SnRK2 via its C-terminal VAST (VAD1 analog of StAR-related lipid transfer) domain with high confidence. BAGP1 is a membrane-associated protein containing both C2 and GRAM domains – features commonly linked to lipid binding. This suggests an integrating membrane dynamics or lipid signaling into osmotic stress responses via SnRK2 signaling.

We identified several conserved candidates in the Co-IP screen (Fig. 3d,e), including PP2Cs as well as ATP-binding cassette subfamily G transporters (ABC-G transporters). ABC-G transport various substrates, including phytohormones such as ABA, and cuticle and sporopollenin precursors, suggesting a potential link to stress response or reproductive development. We also detected WD40-repeat domain proteins, which typically function as scaffolds for protein complex assembly. In *Arabidopsis*, this includes Varicose-Related Protein (VCR), a known target of SnRK2 involved in the formation of the mRNA decapping complex^64^; in *Mesotaenium*, we identified another component of the decapping complex, suggesting a role for SnRK2 in mRNA decapping. Interactors also included pentatricopeptide repeat (PPR) proteins – a large family often associated with RNA processing and organellar gene expression^65^ – and the Calcium-Sensing Receptor (CaS), a chloroplast-localized protein involved in calcium signaling^66^. However, given SnRK2’s expected cytosolic or nuclear localization, this may represent a non-specific interaction. HKs and receptor-like kinases (RLKs) were also recovered, both of which may act as upstream regulators of SnRK2 signaling. Among the RLKs of *Arabidopsis*, we identified Cytokinin Response 1 Kinase (CARK), which has previously been implicated in ABA signal transduction^67,68^. Another putative link to ethylene signaling was the central ethylene signaling protein Ethylene Insensitive 2 (EIN2), identified in algal species but not in the embryophytes (Fig. 3e). Finally, we observed several species-specific interactors, such as the osmotic sensor and calcium influx regulator OSCA in *Zygnema*.

Overall, our systematic cross-species Co-IP analysis of SnRK2 provides a unique comparative framework to investigate the conservation and diversification of SnRK2 signaling across the green lineage (Fig. 3e).

### Divergent and conserved tracks for the role of ABF in hyperosmotic stress response

To classify the *Mesotaenium* ABF, we constructed a phylogenetic tree (Fig. 4a). Rather than analyzing the entire bZIP family – which is too large and diverse for this purpose – we restricted our analysis to a subset of bZIP proteins with high sequence similarity to the *Mesotaenium* ABF. We recovered a well-supported clade that includes all known *Arabidopsis* ABRE-binding bZIP transcription factors, several related proteins, and *Me*ABF (Fig. 4a). We then focused our analysis on the single *Me*ABF, which, importantly, is phosphorylated in *Mesotaenium* upon both hyperosmotic and cold treatment.

**Figure 4.**
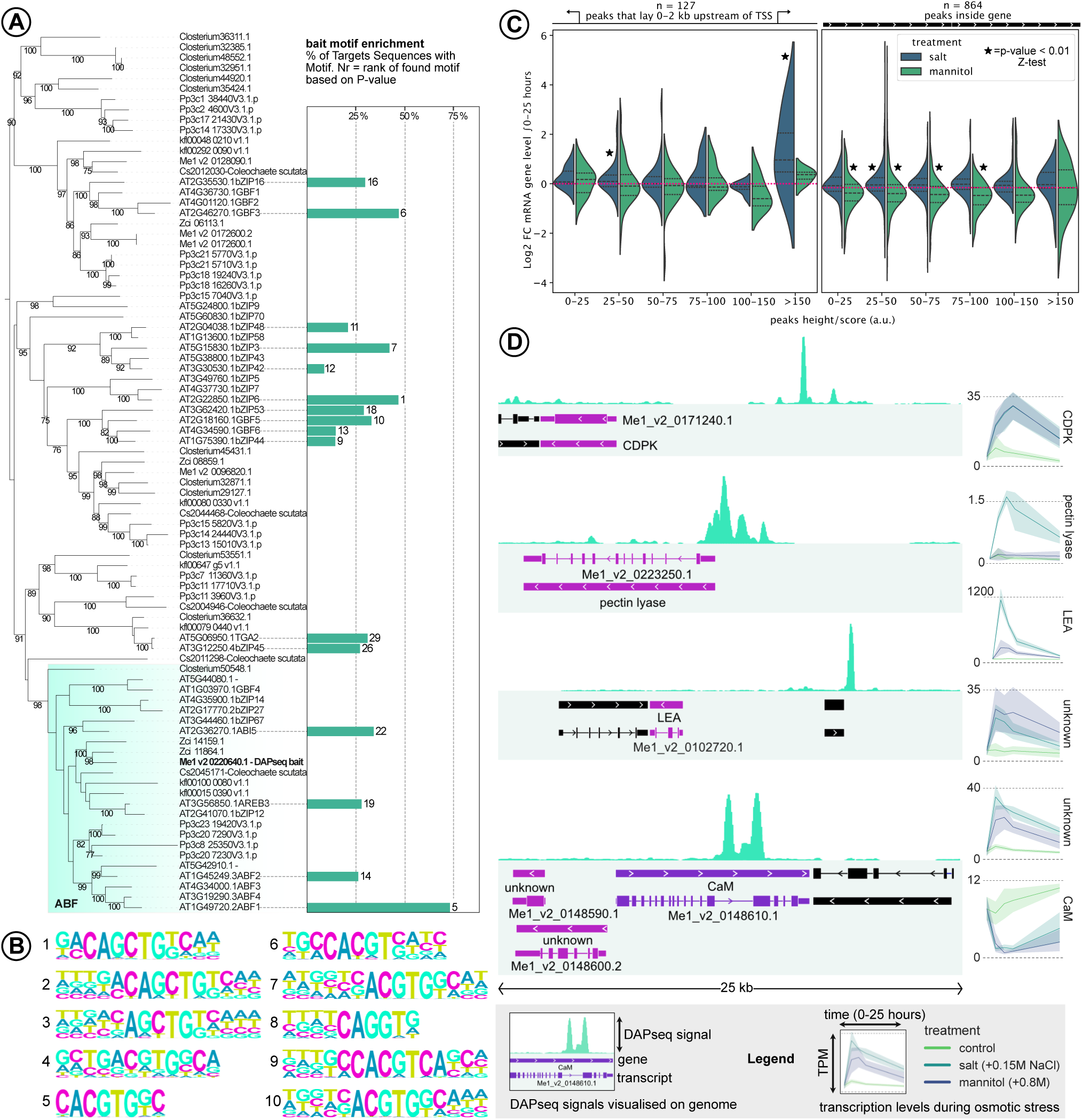
DAP-seq analysis in *Mesotaenium endlicherianum* using ABF as bait. A) Phylogenetic analysis of ABF, with relevant transcription factors from the analysis shown partly in panel B highlighted. Their motif occurrence within the *Mesotaenium* DAPseq analysis and p-value ranking are indicated. B) Top 10 enriched binding motifs (ranked by p-value) identified in ABF-bound regions, corresponding to known motifs of other transcription factors. C) Genomic distribution of ABF binding peaks and associated transcriptional changes under osmotic stress. Left: Genes with putative ABF binding sites within 2 kb upstream of the transcription start site (TSS). Right: Genes with putative binding sites located within gene bodies. Asterisks denote peak groups (based on peak score) whose associated genes exhibit significant changes in expression under hyperosmotic conditions (Z-test). D) Selected high-confidence ABF binding sites (peak height > 200; top 14), shown with their gene neighborhoods. Differentially expressed genes, as reported in Zegers et al.^10^, are indicated.

To characterize the DNA-binding activity of *Me*ABF, we performed DNA Affinity Purification Sequencing (DAP-seq). A total of 3121 ABF binding sites were identified across the genome (FDR < 0.05). Among the 783 high-scoring sites (peak score > 20), 410 sites were located within gene bodies, and at least one binding site was detected in the promoter region (defined as 5 kb upstream) of 758 genes. These genes were subsequently subjected to GO enrichment analysis. Genes containing an ABF binding site within their gene body showed significant enrichment (Bonferroni-corrected p < 0.05) for terms related to DNA repair, protein binding, and the integrator complex. In contrast, genes with a binding site located within 5 kb upstream of their transcription start site (TSS) did not exhibit any significantly enriched GO terms. The enriched peak area sequences showed clear enrichment for several canonical bZIP-binding motifs (Fig. 4b), confirming the success of the assay. Notably, the known binding motif for *Arabidopsis* ABF3 was highly overrepresented, alongside motifs associated with other non-ABF bZIP transcription factors (Fig. 4b).

To assess potential functional relevance, we compared predicted binding sites of the Mesotaenium ABF with gene expression data under hyperosmotic stress conditions, as previously reported^1^. We observed a modest association between the presence of ABF binding peaks – either directly upstream of transcription start sites or within gene bodies – and differential gene expression with several peak groups showing statistically significant enrichment (Z-test, p<0.05). Specifically, under salt stress, genes with detected binding sites in their promoter regions or within their coding sequences tended to be modestly upregulated (mean Log2 fold change = +0.22, p_Z-test_<3E-4). In contrast, mannitol treatment was associated with a slight downregulation (mean Log2 fold change = −0.22, p_Z-test_<8E-18) of genes harboring intragenic binding sites.

We manually examined the top-scoring peaks and their surrounding genomic regions within a 60 kbp window (Fig. 4d and Supplementary Fig. 3). Notably, we identified a broad binding site located in the promoter region of a pectin lyase gene, which is implicated in the salt stress response of *Mesotaenium*. In addition, we detected a putative binding site upstream of both a CDPK gene and a late embryogenesis abundant (LEA) gene. The CDPK gene has here previously been suggested to participate in the ABF signaling cascade, while the LEA gene is a well-established target of ABF transcription factors^69,70^. However, these binding sites lie far from their transcription start sites linearly; yet having no intervening genes, their 3D distance may be quite short. We also identified a potential ABF binding site within a gene encoding calmodulin (CaM), another key component of calcium signaling. Notably, this gene exhibited significant transcriptional repression under hyperosmotic stress conditions.

These findings suggest that ABF plays a regulatory role in the hyperosmotic stress response – particularly in response to salt stress – and possibly linked to a calcium signaling loop in *Mesotaenium*.

### Osmotic stress response in the absence of detectable ABA

The evolutionary origin of phytohormone response cascades is a major question in plant evo-devo research. In the closest algal relatives of land plants, ABA has so far only been detected in certain species and often at miniscule levels^71,72^. This aligns with the PYL homolog of *Zygnema* not binding ABA and inhibiting PP2C in an ABA-independent manner^39^. Yet, ABA might be induced upon hyperosmotic stress response of *Mesotaenium* – and, by extension, in the potential phosphorylation of ABF. To scrutinize this, we employed two complementary approaches: an apocarotenoid screening (Fig. 5a) and a targeted phytohormone analysis (Fig. 5b). In neither assay was ABA detected. Like other streptophyte algae^72,73^, *Mesotaenium* lacks the NCED enzyme^34^, which is essential for ABA biosynthesis via the canonical carotenoid cleavage pathway. While the three cytosolic enzymes of ABA biosynthesis can complement the respective *Arabidopsis* knock-out mutant^74^, in the algal it may be synthesized through an alternative, non-canonical pathways involving other carotenoid cleavage dioxygenases (CCDs)^75^. These enzymes can produce ABA precursors such as Apo11 and OHApo11, or ABA biosynthesis can be induced or repressed by other apocarotenoids as in the case of OHApo13 (zaxinone)^76,77^ and Apo9 (β-ionone^78^), respectively. While we detected no ABA, we successfully detected the majority of other targeted apocarotenoids. Notably, the levels of OHApo11, Apo11, and OHApo13 did not change significantly after 2 or 6 hours of salt treatment. Under cold treatment, these compounds exhibited a modest increase, along with a broader rise in apocarotenoid levels compared to untreated controls – suggesting a general upregulation of carotenoid cleavage in response to low temperatures after few hours which aligns with previous findings in *Mesotaenium*^79^.

**Figure 5.**
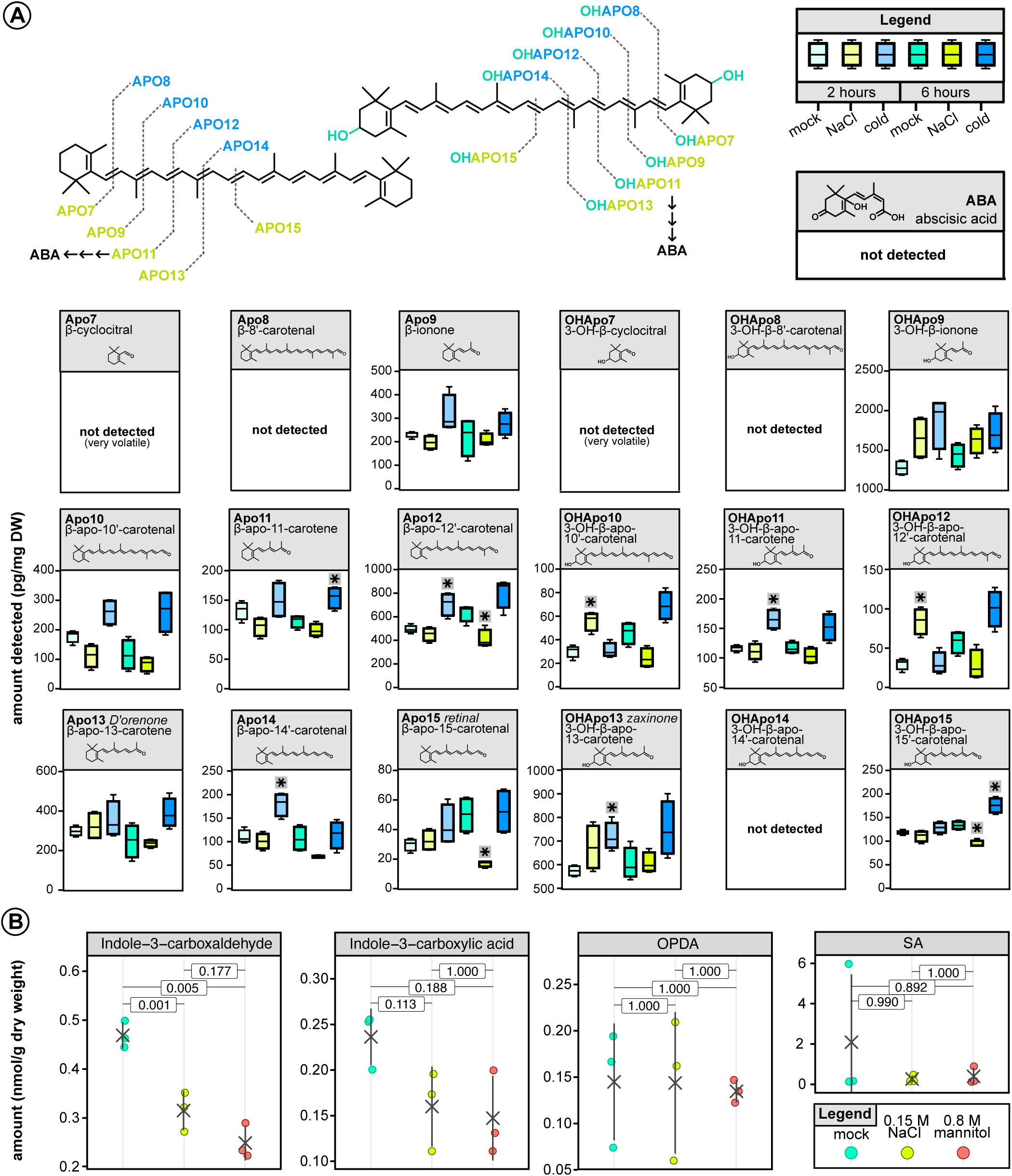
Apocarotenoid and phytohormone levels in *Mesotaenium endlicherianum* under abiotic stress. A) Apocarotenoid concentrations measured after 2 and 6 hours of exposure to stress treatments: 0.15 M NaCl (salt), a cold treatment of –Δ12 °C, and a mock control. Asterisks above the boxplots indicate treatments that differ significantly from the control (p < 0.05; t-test with Benjamini–Hochberg correction for multiple comparisons). B) Phytohormone levels measured after 3 hours of treatment with mock, 0.15 M NaCl, or 0.8 M mannitol. Crosses indicate the mean amount; p-values are displayed above the bars and were Bonferroni-adjusted for multiple comparisons. The apocarotenoid and phytohormone measurements were performed independently and notably, ABA was not detected in either dataset.

In the phytohormone screen (Fig. 5b), only four compounds could be reliably quantified: indole-3-carboxyaldehyde, indole-3-carboxylic acid, 12-oxo-phytodienoic acid (OPDA) and salicylic acid (SA). Of the four detected phytohormones, only indole-3-carboxyaldehyde showed a significant response to osmotic stress: its levels dropped by up to 50% following 3-hour treatments with 0.15 M NaCl and 0.8 M mannitol. This decline may indicate disruptions in tryptophan metabolism, or a potential change in signaling pathways, although these interpretations remain speculative.

Taking previously published work, our work, and these modest changes into account, we would argue that the response to osmotic stress in the closest algal relatives of land plants appears to be driven by other secondary messengers than classical phytohormones known from angiosperms.

### Coalescing tracks in the evolution of the embryophytic stress signaling cascades

Using SnRK2 and ABF as beacons, we investigated their integration into stress response cascades in one of the closest algal relatives of land plants and retraced the evolution of streptophyte signaling systems (Fig. 6). In the LCA of Zygnematophyceae and embryophytes, ABF was regulated by CDPKs in response to calcium influx triggered by hyperosmotic stress. This interaction likely initiated the transcription of stress-responsive genes, including *LEA*. Our data show that *Me*ABF is the most significantly phosphorylated transcription factor in *Mesotaenium* after one hour of hyperosmotic stress and *Me*CDPKs and *Me*ABF likely interact. We also identified putative ABF binding sites upstream of *LEA* and *CDPK* genes that are transcriptionally upregulated under hyperosmotic stress. Supporting evidence from *Arabidopsis thaliana*: (1) hyperosmotic stress induces calcium influxes^33^; (2) calcium activates CDPKs^34^; (3) at least two CDPKs have been shown to phosphorylate and regulate ABFs^17,35^; and (4) ABFs are known to induce *LEA* expression^27,69^. While it remains possible that these components were assembled independently in different lineages, the most parsimonious explanation is that the CDPK-ABF signaling axis was already present in their shared ancestor; this aligns with their co-expression being predicted for the LCA of land plants and zygnematophytes^38^. This proposed incorporation of CDPK into part of the ABA signaling pathway in the LCA is further supported by a regulatory network inference across nine species, including three algae, which predicted this interaction^38^.

**Figure 6.**
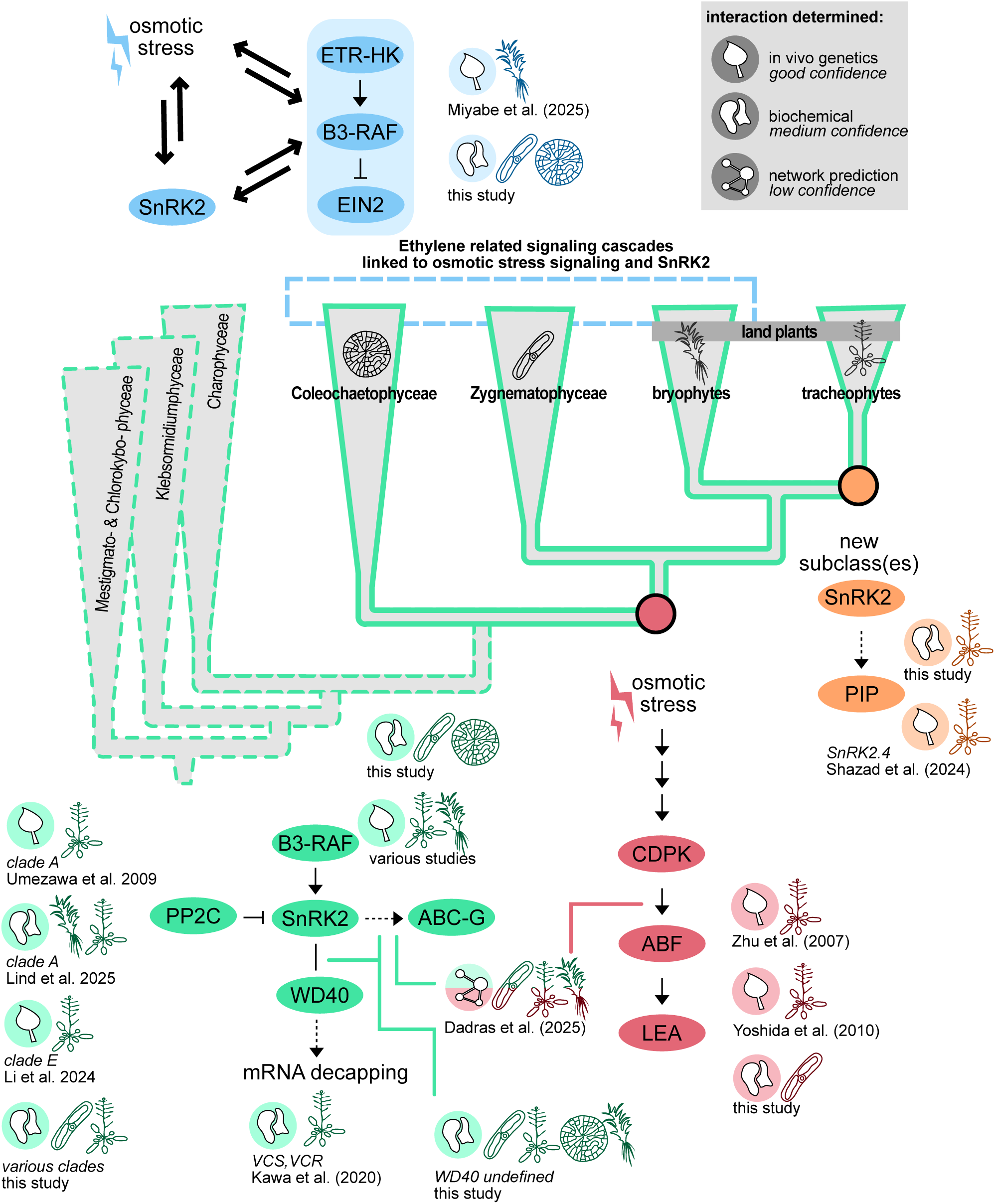
Evolutionary scenarios for the assembly of the osmotic stress signaling network. Work from this and other studies synthesized into complementary models that brought together PP2C, SnRK2, and ABF during streptophyte evolution.

In the LCA of Coleochaetophyceae and embryophytes, SnRK2 interacted with a conserved set of proteins, including B3-type RAF kinases, WD40-domain proteins, PP2Cs, and ABC-G transporters. Our evidence for PP2C-SnRK2 interaction is currently limited to Zygnematophyceae and embryophytes, and the precise identity of the ancestral PP2C clade involved remains unresolved. PP2CA stands out as the most likely candidate, given its highly conserved interaction with SnRK2 across embryophytes^80^. However, direct functional evidence is still lacking. Notably, SnRK2 from *Klebsormidium* can be inhibited by *Arabidopsi*s PP2CA^29^, but the capacity of algal PP2Cs to inhibit SnRK2 has yet to be tested. The absence of PP2CA in both the Y2H screen and the Co-IP experiments does not exclude the possibility of interaction in the algal context. One explanation is that PP2CA is present at relatively low abundance compared to SnRK2 (Supplementary Fig. 4), reducing its likelihood of detection in these assays. In addition, the experimental conditions or protein conformations may not have been conducive to stable binding, further limiting the detection of potential interactions. The WD40-domain proteins and ABC-G transporters emerge as promising conserved SnRK2 interactors, yet both families are large and functionally diverse. WD40 proteins are known to act as scaffolds, particularly in mRNA degradation pathways. For example, the VARICOSE family of WD40 proteins, known interactors of SnRK2^23^, function as scaffolds in mRNA decapping, thereby regulating mRNA degradation. In *Mesotaenium*, we identified a WD40-domain protein with homology to a component of the ubiquitination complex, suggesting a possible link between SnRK2 and regulated protein turnover. Growing evidence from *Arabidopsis*^81^ indicates that B3-RAF kinases act as upstream activators of SnRK2. Our results extend this model into the algal lineage: we found moderate evidence for a B3-RAF-SnRK2 interaction in *Mesotaenium* and weaker – but still noteworthy – evidence in *Coleochaete*. Given the scarcity of molecular data from Coleochaetophyceae, even preliminary interaction evidence is valuable and contributes to our understanding of early signaling evolution. What lies upstream of B3-RAF kinases^31^ in algae remains unknown. By analogy with embryophytes, where HKs are candidate upstream regulators of B3-RAFs^82^, a similar mechanism may have existed in ancestral streptophytes.

Miyabe et al.^83^ proposed an evolutionary model in which ethylene and SnRK2 signaling were originally integrated, and only later during tracheophyte evolution did they evolve into distinct, independent pathways. This model is particularly compelling for *Physcomitrium*, which possesses only a single B3-type RAF kinase. Given that B3-RAF kinases function upstream of SnRK2 and are also known to act as central components in ethylene signaling, it is reasonable to infer that these pathways are at least to some extent integrated in *Physcomitrium*. Our data extend this concept to Zygnematophyceae and Coleochaetophyceae. We detected EIN2 – an essential downstream effector of the ethylene signaling pathway^84,85^ – in the Co-IPs of SnRK2 in *Coleochaete*, *Mesotaenium*, and *Closterium*, suggesting a possible physical or functional link between the two pathways in this grade. Additionally, several histidine kinases related to ETR proteins, though not the receptors themselves, were phosphorylated or dephosphorylated in response to hyperosmotic stress in *Mesotaenium*. Taken together, these findings offer preliminary support for the hypothesis that ethylene and SnRK2 signaling were interconnected in the last common ancestor (LCA) of phragmoplastophytes. This ancestral linkage may have been gradually uncoupled in vascular plant lineages, leading to the distinct pathways observed in extant tracheophytes.

In tracheophytes (vascular plants), new subgroup(s) of SnRK2 kinases emerged. Undergoing massive subfunctionalization and neofunctionalization, one potential innovation may involve the regulation of plasma membrane intrinsic proteins (PIPs), a subgroup of aquaporins. In our SnRK2 co-immunoprecipitation (Co-IP) experiments in *Arabidopsis*, we identified a striking enrichment of PIPs – an observation not mirrored in any of the algal datasets. Notably, PIP regulation has previously been associated with SnRK2.4^86^, a member of subclass I SnRK2s, which shows less similarity to the algal SnRK2s than does subclass III. This suggests that the interaction between SnRK2 and PIPs may represent a tracheophyte-specific innovation. The functional implications of this interaction could be significant. PIPs play a crucial role in facilitating lateral water transport within the vascular system and are also implicated in hydrogen peroxide (H₂O₂) signaling. The emergence of SnRK2-mediated regulation of PIPs may have been an important adaptation for managing water transport and redox signaling in vascular plants. Further research is needed to clarify the role of SnRK2-PIP interactions in the context of vascular function, and we encourage future studies to explore this potential link in greater detail.

## Conclusion

Our integrative analysis reveals that key components of the canonical ABA signaling cascade were assembled from pre-existing and partially independent signaling modules in a streptophyte algal ancestor of embryophytes. By combining phosphoproteomics, protein-protein interaction mapping, and genome-wide transcription factor binding analyses, we propose a model that ABF functions as a central regulator of osmotic stress responses in *Mesotaenium* despite the absence of detectable ABA. This highlights that core transcriptional outputs of the ABA pathway predate the hormone itself and were likely driven by alternative upstream signals, particularly calcium-dependent pathways mediated by CDPKs. In contrast, SnRK2 appears to occupy a more complex and potentially context-dependent role, interacting with conserved partners such as RAF kinases and PP2Cs while also linking to novel regulatory layers including translation, membrane dynamics, and mRNA turnover. The strong involvement of histidine kinases and the observed connections to ethylene signaling further suggest that ancestral stress response networks were highly interconnected. Together, our findings support a model in which modern ABA signaling emerged through the coalescence of ancient, parallel signaling circuits — prior to plant terrestrialization.

## METHODS

### Phosphoproteomics

The alga *Mesotaenium endlicherianum* SAG12.97 was sourced for this and all following experiments from the Culture Collection of Algae at Göttingen University (SAG)^87^. The *Mesotaenium* gene model V2, produced by Dadras et al.^88^, was used for this and all subsequent experiments. *Mesotaenium* was cultured on cellophane placed over 1.5% agar-solidified C-medium^89^ in 14 cm diameter plates under a 16-hour light (∼*40* µE)/8-hour dark cycle at 18 °C. After 8 days of growth, 3 hours (±2.5 minutes) after the grow lights turned on, the cellophane with the attached algae was transferred to new plates under sterile conditions. The treatments included a control with no supplementation, a salt treatment with 150 mM NaCl, a high osmotic treatment with 800 mM mannitol, and a cold treatment in which plates were placed on ice. For the cold treatment, the temperature was monitored and it decreased to 6.3 °C (±0.3 °C) over 15 minutes after which it stabilised. Algal samples were collected after 1 and 3 hours by scraping the biomass of three plates of the same condition into Eppendorf tubes, which were immediately flash-frozen in liquid nitrogen. The stress treatment was repeated for 5 biological replicates.

From the flash-frozen samples, total proteome and phosphopeptide-enriched samples were prepared. The frozen tissue was grinded with 2 2.5mm steel beads in a Retsch mill, then 300 µL extraction buffer (8M urea, 20 µL/mL Phosphatase Inhibitor Cocktail 2 (Sigma, P5726-5ML), 20 µL/mL Phosphatase Inhibitor Cocktail 3 (Sigma, P0044-5ML),5 mM DTT) was added and samples were incubated for 30 min with shaking, after which cell debris was removed by centrifugation. Samples were alkylated with CAA (550 mM stock, 14 mM final), the reaction was quenched with DTT (5 mM final). Next, samples were diluted to 4M urea with 100 mM Tris-HCl pH 8.5, 1 mM CaCl_2_ and digested with 5 µg LysC (stock: 1 µg/µL Lys-C (Merck) in H_2_O) for 3h at RT. Subsequently, samples were diluted with 100 mM Tris-HCl pH 8.5, 1 mM CaCl_2_ to 1M urea, then 5 µg trypsine (stock: 1 µg/µL in 1 mM HCl,) was added and samples were, the samples were mixed and incubated o/N at 37 °C. After incubation, samples were was acidified with TFA to 0.5% final concentration and samples were desalted using C18 SepPaks (1cc cartridge, 100 mg (WAT023590)). In brief, SepPaks were conditioned using methanol (1 mL), buffer B (80% acetonitrile, 0.1% TFA) (1 mL) and buffer A (water, 0.1% TFA) (2 mL). Samples were loaded by gravity flow, washed with buffer A (1 x 1 mL, 1x 2 mL) and eluted with buffer B (2 x 400 µL). 42 µL of eluates were used for peptide measurement and total proteome analysis.

For the library samples, 2 µL aliquots from all samples were mixed and submitted to SCX fractionation. To this end, StageTips were prepared using 6 layers of SPE disk (Empore Cation 2251 material) activated with acetonitrile and washed with 1% TFA (100 µL each) and buffer A (water, 0.2% TFA) (100 µL) by spinning 5 min (1.5k x g), samples were acidified to 1 % TFA, loaded by centrifugation (10 min, 800 x g) and washed with buffer A (5 min, 1.5k x g) (100 µL). Fractionation was carried out using an ammonium acetate gradient (20% ACN, 0.5 % FA) starting from 25 mM to 500 mM for 9 fractions and two final elution steps using 1% ammonium hydroxide, 80% ACN and finally 5% ammonium hydroxide, 80% ACN. All fractions were eluted by centrifugation (1 min, 1200 x g) using 2 x 30 µL eluent. The fractions were dried and taken up in 10 µL A* buffer. Peptide concentration was determined by Nanodrop and samples were diluted to 0.2 µg/µL for measurement.

For total proteome analysis, the remaining 40 µL eluted peptides from the SepPack purification were dried and then taken up in 10 µL A* buffer. Peptide concentration was determined by Nanodrop and samples were diluted to 0.2 µg/µL for measurement.

For phosphopeptide enrichment by metal-oxide chromatography (MOC) (adapted from: Nakagami^90^) the remaining samples were evaporated to a sample volume of 50 µL and diluted with sample buffer (2 mL AcN, 820 µL lactic acid (LA), 2.5 µL TFA / 80% ACN, 0.1% TFA, 300mg/ml LA, final concentrations) (282 µL). Since all samples showed a small pellet after dilution, the samples were centrifuged briefly and only the supernatant was used for phosphopeptide enrichment. MOC tips were prepared by loading a slurry of 3mg/sample TiO_2_ beads (Titansphere TiO_2_ beads 10 µm (GL Science Inc, Japan, Cat. No. 5020-75010)) in 100 µL MeOH onto a C2 micro column and centrifugation for 5 min at 1500g. Tips were washed with centrifugation at 1500g for 5 min using 75µL of solution B (80% acetonitrile, 0.1% TFA) and 75 µL of solution C (300 mg/mL LA in solution B). To simplify the processing, samples tips were fitted onto a 96/500 µL deep well plate (Protein LoBind, (Eppendorf Cat. No. 0030504100). After washing MOC tips were transferred to a fresh plate, samples were loaded onto the equilibrated tips and centrifuged for 10 min at 1000g. The flow through was reloaded onto the tips and centrifugation was repeated. Tips were washed with centrifugation at 1500g for 5 min using 75 µL of solution C and 3x 75µL of solution B. For the elution of the enriched phosphopeptides the tips were transferred to a fresh 96/500 µL deep well plate containing 100 µL/well of acidification buffer (20% phosphoric acid). Peptides were eluted first with 50 µL elution buffer 1 (5% NH_4_OH) and centrifugation for 5 min at 800g, then with 50 µL of elution buffer 2 (10% piperidine) and centrifugation for 5 min at 800g. Next, the samples were desalted using StageTips with C18 Empore disk membranes (3 M) (Rappsilber et. al., Anal. Chem. 2003, 75, 663.), dried in a vacuum evaporator, and dissolved in 10 µL 2% ACN, 0.1% TFA (A* buffer) for MS analysis.

Total proteome and phosphoproteome samples were analyzed using LC-MS/MS with data-dependent acquisition. All samples were analyzed using an Ultimate 3000 RSLC nano (Thermo Fisher) coupled to an Orbitrap Exploris 480 mass spectrometer equipped with a FAIMS Pro interface for Field asymmetric ion mobility separation (Thermo Fisher). Peptides were pre-concentrated on an Acclaim PepMap 100 pre-column (75 µM x 2 cm, C18, 3 µM, 100 Å, Thermo Fisher) using the loading pump and buffer A with a flow of 7 µl/min for 5 min. Peptides were separated on 16 cm frit-less silica emitters (New Objective, 75 µm inner diameter), packed in-house with reversed-phase ReproSil-Pur C18 AQ 1.9 µm resin (Dr. Maisch). Peptides were loaded on the column and eluted for 130 min using a segmented linear gradient of 5% to 95% solvent B (0 min : 5%B; 0-5 min −> 5%B; 5-65 min −> 20%B; 65-90 min −>35%B; 90-100 min −> 55%; 100-105 min −>95%, 105-115 min −>95%, 115-115.1 min −> 5%, 115.1-130 min −>5%) (solvent A 0% ACN, 0.1% FA; solvent B 80% ACN, 0.1%FA) at a flow rate of 300 nL/min. For the total proteome and fractionated (library samples), mass spectra were acquired in data-dependent acquisition mode with a TOP_S method using a cycle time of 2 seconds. For Field asymmetric ion mobility separation (FAIMS) two compensation voltages (−45 and −60) were applied, the cycle time for each experiment was set to 1 second. MS spectra were acquired in the Orbitrap analyzer with a mass range of 320-1200 m/z at a resolution of 60,000 FWHM and a normalized AGC target of 300%. Precursors were filtered using the MIPS option (MIPS mode = peptide), the intensity threshold was set to 5000, Precursors were selected with an isolation window of 1.6 m/z. HCD fragmentation was performed at a normalized collision energy of 30%. MS/MS spectra were acquired with a target value of 75% ions at a resolution of 15,000 FWHM, at an automated injection time and a fixed first mass of m/z 100. Peptides with a charge of +1, greater than 6, or with unassigned charge state were excluded from fragmentation for MS2, dynamic exclusion for 40s prevented repeated selection of precursors.

For the phosphoproteome analysis, mass spectra were acquired in data-dependent acquisition mode with a TOP_S method using a cycle time of 2 seconds. For field asymmetric ion mobility separation (FAIMS) two compensation voltages (−45 and −65) were applied, the cycle time for the CV-45 experiment was set to 1.2 seconds and for the CV-65 experiment to 0.8 sec. MS spectra were acquired in the Orbitrap analyzer with a mass range of 320-1200 m/z at a resolution of 60,000 FWHM and a normalized AGC target of 300%. Precursors were filtered using the MIPS option (MIPS mode = peptide), the intensity threshold was set to 5000, Precursors were selected with an isolation window of 1.6 m/z. HCD fragmentation was performed at a normalized collision energy of 30%. MS/MS spectra were acquired with a target value of 75% ions at a resolution of 15,000 FWHM, at an injection time of 120 ms and a fixed first mass of m/z 120. Peptides with a charge of +1, greater than 6, or with unassigned charge state were excluded from fragmentation for MS2.

The raw data from to total proteome were processed using MaxQuant software with label-free quantification (LFQ) and iBAQ enabled (Cox et al., Nat. Protoc. 2016, 11, 2301.). Library samples and DDA samples were grouped into separate parameter groups. In the group specific parameters, in the Misc. setting the Match type for library samples was set to “match from” and for DDA to “match from and to”.

MS/MS spectra were searched by the Andromeda search engine against a combined database containing the sequences from *Mesotaenium* and sequences of 248 common contaminant proteins and decoy sequences. Trypsin specificity was required and a maximum of two missed cleavages allowed. Minimal peptide length was set to seven amino acids. Carbamidomethylation of cysteine residues was set as fixed, oxidation of methionine and protein N-terminal acetylation as variable modifications. The match between runs option was enabled. Peptide-spectrum-matches and proteins were retained if they were below a false discovery rate of 1% in both cases.

Statistical analysis of the MaxLFQ values was carried out using Perseus. Quantified proteins were filtered for reverse hits and hits “only identified by site” and MaxLFQ values were Log2 transformed and samples were grouped by condition. Next, the data was separated for a mixed imputation processing: hits were filtered for 3 valid values in one of the conditions. Then, the data was separated into two sets: one set containing mostly missing at random (MAR) hits and the other set containing mostly missing not at random (MNAR) hits by filtering the data for 1 valid hit in each group and splitting the resulting matrices^91^. The resulting matrix with at least 1 valid hit in each group is the MAR dataset, the matrix with the hits filtered out is the MNAR dataset. The missing values of each dataset were then imputed using different options of the “imputeLCMD” R package: the missing values from the MAR dataset were imputed using a nearest neighbor approach (KNN, n=4), the missing values from the MNAR dataset were imputed using the MinProb option (q=0.01. tune.sigma=1). After merging of the imputed datasets two-sample Student’s t-tests were performed using a permutation-based FDR of 5%. Alternatively, volcano plots were generated using an FDR=0.05 and an S0=1. The Perseus output was exported and further processed using Excel.

Also for the data analysis of the phosphoproteomics, raw data were processed using MaxQuant^92^. MS/MS spectra were searched by the Andromeda search engine against a combined database containing the sequences from *Mesotaenium* and sequences of 248 common contaminant proteins and decoy sequences. Trypsin specificity was required and a maximum of two missed cleavages allowed. Minimal peptide length was set to seven amino acids. Carbamidomethylation of cysteine residues was set as fixed, phosphorylation of serine, threonine and tyrosine, oxidation of methionine and protein N-terminal acetylation as variable modifications. The match between runs option was enabled. Peptide-spectrum-matches and proteins were retained if they were below a false discovery rate of 1% in both cases.

Statistical analysis was carried out on phospho peptide level using the intensities obtained from the “modificationSpecificPeptides” output. Using these data, missing values were imputed using random sampling from normal distributions whose parameters depended on the degree of missingness within each condition. This approach assumes the following: if a phosphopeptide is not detected in any replicate of a given condition, it is likely unphosphorylated in that condition. Conversely, if a phosphopeptide is detected in some but not all replicates of a condition, it is interpreted as being present but occasionally below the detection limit. Samples from the 1-hour and 3-hour treatments were analyzed separately, generating two independent datasets. Each dataset consisted of four experimental conditions with five replicates each (20 samples in total). The imputation procedures are as followed:

- Imputation method A) When 1,2,3, or 4 out of 5 replicate values were missing, missing data were imputed by sampling from a normal distribution with a standard deviation equal to 0.5 × the standard deviation of all observed values within the corresponding sample. The mean (μ) of this distribution was set to the median of all observed values from peptides of the same missing-value category, shifted downward by λ(z⍰) × σ, where σ is the standard deviation of the sample and λ(z⍰) = φ(z⍰)/(1 - Φ(z⍰)) is the inverse Mills ratio. Here, z⍰ is defined such that 1 - Φ(z⍰) = p, where p ∈ [0.2, 0.4, 0.6, 0.8] corresponds to the proportion of missing values (4, 3, 2, or 1 missing replicates, respectively), so that λ(z⍰) reflects the expected standardized shift for the corresponding upper-tail truncation.
- Imputation method B) When all 5 values were missing, the normal distribution parameters σ was based on all measured intensities for the same peptide across other conditions. The distribution mean µ was set to the mean of all detected peptides in the corresponding sample and then downshifted by 5 times the standard deviation of those detected peptides in the corresponding sample.

The observed and imputed data obtained from Method A followed a normal distribution (see Supplementary Fig. 1), allowing for the calculation of ANOVA p-values for all phosphopeptides for which all conditions contained at least 1 observed value. P-values were corrected for multiple testing using the false_discovery_control function of the scipy.stats package. To be able to also include peptides with conditions containing entirely missing values, two heuristic reliability metrics were developed: the Reliability-Adjusted F-score (RAF) for multiple-condition comparisons and the Reliability-Adjusted T-score (RAT) for pairwise comparisons. These metrics help identify phosphopeptides likely to differ between conditions but do not represent formal test statistics. They were calculated as follows:

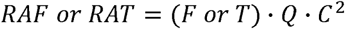

where:

- F and T represent the standard ANOVA F-statistic and the Student’s t-statistic, respectively.
- Q is the proportion of observed (non-imputed) data across all samples. This term reduces the score linearly with increasing missingness.
- C is the bimodal polarization index, which penalizes uneven distributions of missing values across conditions. To compute C, the number of missing values per condition is determined. In the case of four conditions, C is computed as follows: *C* = (*n*_1_ + (0.5 · *n*_2_))/4, where n₁ is the count of the most frequent number of missing values and n₂ is the count of the second most frequent. In the case of pairwise comparisons, C is set to 1.

For ease of data interpertation, the RAF and RAT scores were transformed into a surrogate −log p-values using the python expression “−math.log(1 - stats.f.cdf(RAF, 3, 16))”, where the RAF score is treated as an F-statistic with 3 and 16 degrees of freedom and the excel formula “LN(T.DIST.2T(RAT, 8))”, where the RAT score is treated as an t-statistic and 8 are the degrees of freedom”. Likewise, the adjusted ANOVA p-values were log-transformed using the Python expression −math.log(p).

All data points were visualized using Plotly within a Shiny app (Supplementary Data 2 or https://jaccoline.github.io/Mesotaenium-Phosphoproteomics-DataExploration/). Functional annotation included the top BLASTp hits to Arabidopsis thaliana proteins and EggNOG annotations from Dadras et al.^88^ Peptides were classified as belonging to protein kinases if their protein group contained InterPro domains IPR011009 or IPR005467. Transcription-associated proteins (TAPs) were identified based on the TAPscan v4 database^47^, except for certain histidine kinases, which were already categorized as protein kinases.

### Hormonal measurements

*Mesotaenium* was cultured on cellophane placed over 1.5% agar-solidified C-medium in 14 cm diameter plates under a 16-hour light (∼70 µE)/8-hour dark cycle at 18 °C. After 13 days of growth, 2 hours (±5 minutes) after the grow lights turned on, the cellophane with the attached algae was transferred to new plates under sterile conditions. The treatments included a control with no supplementation, a salt treatment with 150 mM NaCl, and a high osmotic treatment with 800 mM mannitol. Algal samples were collected after 3 hours by scraping the biomass of three plates of the same condition into 2 mL Eppendorf tubes, which were immediately flash-frozen in liquid nitrogen. The stress treatment was repeated for 3 biological replicates. The samples were then lyophilized using a VaCo 2 Zirbus lyophilizer.

Extraction and LC/tandem MS analysis was performed as described^93^. 10 mg homogenized and lyophilized plant material was extracted together with an internal standard-mix containing 10 ng D_4_-SA (C/D/N Isotopes Inc., Pointe-Claire, Canada), 30 ng D_5_-oPDA (kindly provided by Otto Miersch, Halle/Saale, Germany), 10 ng D_6_-ABA (C/D/N Isotopes Inc., Pointe-Claire, Canada), 20 ng D_5_-IAA (Eurisotop, Freisig, Germany).

Reversed phase separation of constituents was achieved by using an ACQUITY UPLC® system (Waters Corp., Milford, MA, USA) and analysed by nanoelectrospray ionization (nanoESI) (TriVersa Nanomate^®^; Advion BioSciences, Ithaca, NY, USA) coupled with an AB Sciex 4000 QTRAP^®^ tandem mass spectrometer (AB Sciex, Framingham, MA, USA) employed in scheduled multiple reaction monitoring mode. Mass transitions corresponding to 63 phytohormones and related species were investigated (see Supplementary Data 6), but only four were reliably detected. Their mass transitions and the one for ABA were as follows: 137/93 [declustering potential (DP) −25 V, entrance potential (EP) −6 V, collision energy (CE) −20 V] for SA, 141/97 (DP −25 V, EP −6 V, CE −22 V) for D_4_-SA, 296/170.2 (DP −65 V, EP −4 V, CE −28 V) for D_5_-oPDA, 291/165 (DP −50 V, EP −5 V, CE −26 V) for oPDA, 179/135 (DP −35 V, EP −9 V, CE −14 V) for D_5_-IAA, 160/116 (DP −40 V, EP −6.5 V, CE −22 V) for indole-3-carboxylic acid, 269/159 (DP −30 V, EP −5 V, CE −16 V) for D_6_-ABA, 263/153 (DP −35 V, EP −4 V, CE −14 V) for ABA in negative ionization mode, and 181/134 (DP 41 V, EP 10 V, CE 21 V) for D_5_-IAA and 146/118 (DP 51 V, EP 3.5 V, CE 19 V) for indole-3-carboxaldehyde in positive ionization mode. Quantification was carried out using a calibration curve of intensity (*m/z*) ratios of [unlabeled]/[deuterium-labeled] *vs.* molar amounts of unlabeled (0.3-1000 pmol). P-values were calculated and adjusted using the R package ggpmisc and function stat_multcomp with p.adjust.method set to Bonferroni.

#### Apocarotenoids profiling by LC-MS/MS analysis

*Mesotaenium* was cultured on cellophane placed over 1.5% agar-solidified C-medium in 14 cm diameter plates under a 16-hour light (∼80 µE)/8-hour dark cycle at 20 °C. After 8 days of growth, 3 hours (±5 minutes) after the grow lights turned on, the cellophane with the attached algae was transferred to new plates under sterile conditions. The treatments included a control with no supplementation, a salt treatment with 150 mM NaCl, and a cold treatment in which plates were placed on ice without additional supplementation. For the cold treatment, the temperature was monitored and it decreased to 8 °C over 15 minutes after which it stabilised. The initiation of stress treatments required transferring the samples from the growth chamber to a sterile bench and back, resulting in a 20-minute period of fluctuating light conditions. Algal samples were collected after 2 and 6 hours by scraping or pouring the biomass into 2 mL Eppendorf tubes, which were immediately flash-frozen in liquid nitrogen. The stress treatment was repeated for 6 biological replicates. The samples were then lyophilized using a VaCo 2 Zirbus lyophilizer and ground with a Retsch mill at 20 Hz for 1 minute. The resulting dry weight was adjusted to 25 mg (±1 mg).

Aliquots (25 mg) were transferred to 4-mL amber glass tubes and extracted twice with 1.8 mL of methanol containing 0.1% (w/v) butylated hydroxytoluene (BHT) and spiked with deuterium-labeled apocarotenoid internal standards^94^ (2 ng per sample). Samples were vortexted for 2 min, kept in an orbital shaker at 4 °C for 10 min and sonicated in ice-cooled ultrasound bath for 10 min^94^. After centrifugation for 5 min at 14000 rpm at 4 °C, the combined supernatants were concentrated to dryness under nitrogen gas. Dried extracts were reconstituted in 0.15 mL of acetonitrile, vortexed thoroughly and filtered through a 0.22 µm PTFE filters prior to analysis^95^. LC-MS/MS analysis was performed using a Dionex Ultimate 3000 UHPLC system coupled to a an Orbitrap mass spectrometer (Q Exactive Plus, Thermo Fisher Scientific) equipped with a heated electrospray ionization source. Chromatographic separation was achieved on an ACQUITY UPLC BEH C18 Column (2.1 X 100 mm, 1.7 µm; Water) maintained at 35 °C. The mobile phases consisted of water: acetonitrile/formic acid (80/ 20/ 0.1, v/v/v, solvent A) and acetonitrile/isopropanol/formic acid (60/ 40/ 0.1, v/v/v, solvent B). Separation was carried out using the following gradient: 0-1 min, 100%A; 1-3 min, 100 A to 60%A; 3-8 min, 60%A to 20%A, 8-14 min 20%A to 10%A, 14–15 min, 10%A to 0%A; followed by washing with 100%B and equilibration with 100% A. The flow rate was maintained at 0.2 mL/min and the injection volume was 5 µL. Mass spectrometry analysis was performed in positive ionization mode using parallel reaction monitoring^94,95^ (PRM). Apocarotenoids were identified based on accurate mass measurements, retention times and MS/MS fragmentation spectra in comparison with authentic and isotope labeled standards. Peak integration and data processing were conducted using Thermo Xcalibur 4.8 software. P-values were calculated by comparing the salt and cold treatments to the control using a two-tailed equal-variance *t*-test, and all p-values were subsequently corrected for multiple comparisons using scipy.stats.false_discovery_control.

### DAPseq with ABF

The DAP-seq assay was largely done according to an established protocol with several modifications^96^. In short, genomic DNA was extracted using a phenol:chloroform:isoamylalcohol procedure, followed by a subsequent fragmentation of 5 µg of gDNA into 200-300 bp fragments. Sonicated fragments were end-repaired and Illumina TruSeq adaptors were attached. Next to that, the coding sequence of Me1_v2_0220640.1 was N-terminally fused with a Strep-II tag and expressed using an in vitro wheat germ system, the TNT SP6 High Yield Wheat Germ Mastermix (Promega). Strep-ABF protein was immobilized on Streptavidin beads (IBA), washed 3 times and incubated for 2 hours with 100 ng of gDNA library. After incubation, the beads were washed 3 times with a solution of TBS+0.05% (v/v) IGEPAL. After bead washing, the DNA was eluted and amplified with indexed TruSeq primers. Sequencing was performed on an Illumina Novoseq 6000 platform with 150-bp PE reads by Novogene GmbH. As a control, a biotin-only probe was used for the DAP-seq analysis. Obtained reads were mapped to the *Mesotaenium endlicherianum* SAG12.97 V1 genome^34^. Reads were mapped using Bowtie2 and peaks were called using MACS2, whereby the biotin samples were used as control and the FDR (Benjamini-Hochberg procedure) was set to 0.05. Motif enrichment was performed using the Homer package. Peaks identified using MACS2 (see Supplementary Data 4) were filtered for a minimum peak score of 20, yielding 783 binding sites. Genomic locations were then compared with version 2 of the gene models^88^. Functional enrichment analysis using GO terms and the program FUNC-E was performed on (I) genes containing peaks and (II) genes with binding sites located within 5,000 bp upstream of their transcription start sites.

### Yeast Two-Hybrid Screen

Yeast two-hybrid screening was performed by Hybrigenics Services, S.A.S., Evry, France (http://www.hybrigenics-services.com). The coding sequences for *Me*SnRK2 (geneID: Me1_v2_0141910.1) and *Me*ABF (geneID: Me1_v2_0220640.1) were PCR-amplified and cloned into pB27 as a C-terminal fusion to LexA (LexA-*Me*SnRK2, LexA-*Me*ABF). The constructs were checked by sequencing the entire inserts and each seperately used as a bait to screen a random-primed *M. endlicherianum* SAG12.97 vegative cells cDNA library constructed into pP6. pB27 and pP6 derive from the original pBTM116 plasmid^97^ and pGADGH plasmid^98^, respectively.

A total of 67.8 million clones for the SnRK2 screen and 111 million clones for the ABF screen were screened, corresponding to 6.8 and 11 times the complexity of the screened library (library size ≈ 10 million independent fragments), using a mating approach with YHGX13 (Y187 ade2-101::loxP-kanMX-loxP, matα) and L40ΔGal4 (mata) yeast strains as previously described^99^. Subsequently, 332 (SnRK2-screen) and 91(ABF-screen) His+ colonies were selected on a medium lacking tryptophan, leucine and histidine, and for the SnRK2-screen only supplemented with 0.5 mM 3-aminotriazole to handle bait autoactivation. The prey fragments of the positive clones were amplified by PCR and sequenced at their 5’ and 3’ junctions (see Supplementary Data 5). The resulting sequences were matched with the genes from *M. endlicherianum* SAG12.97 annotation version 2 gene model^88^. A confidence score (PBS, for Predicted Biological Score) was attributed to each interaction as previously described^100^. The PBS relies on two different levels of analysis. Firstly, a local score takes into account the redundancy and independency of prey fragments, as well as the distribution of reading frames and stop codons in overlapping fragments. Secondly, a global score takes into account the interactions found in all the screens performed at Hybrigenics using the same library. This global score represents the probability of an interaction being nonspecific. For practical use, the scores were divided into four categories: “moderate”, “good”, “high” and “very high”. The PBS scores have been shown to positively correlate with the biological significance of interactions^101,102^.

### SnRK2 co-immunoprecipitation

Circa 200 mg (+/− 38 mg) aliquots of wet biomass from leaves from *A. thaliana* Col-0, protonema cultures from *P. patens* (Reute and Gransden), agar-grown culture *M. endlicherianum* SAG 12.97, agar-grown culture *Z. circumcarinatum* SAG 698-1b, liquid-grown culture *Closterium sp.* NIES 68, and liquid-grown *C. scutata* SAG 110.80 were collected in microcentrifuge tubes and immediately frozen in liquid nitrogen. Different leaves, plates, or flasks were used for each biological replicate, respectively. Using liquid nitrogen-cooled 2 ml tube adapters, cells were lysed by bead beating with a Retsch Mixer Mill MM 400 (30 Hz for 30 s intervals) until pulverized. Immediately afterwards ice-cold extraction buffer (50 nM Tris pH 7.5, 150 nM NaCl, 10% Glycerol, 2 mM EDTA, 5 nM DTT, 1% Triton X-100, and one tablet of cOmplete^TM^ ULTRA Tablets Mini, EDTA-free protease inhibitor cocktail (Roche, Grenzach-Wyhlen, Germany, Cat. No. 05892791001) was added, then vortexed. After 30 min incubation on ice, samples were centrifuged (5 min, 4°C, 10,000 g). Supernatants were transferred to a new tube and recentrifuged at (5 min, 4°C, 13,000 g), repeating until no pellet remained. Protein concentration was determined using Pierce™ 660 nm Protein Assay Kit and Reagent (Thermo Fischer, Cat. No.22662). The total protein concentration of the cell lysate was diluted to 0.5 mg/mL. 40 µg of total protein from biological replicates per species was aliquoted for downstream gel plot analysis.

Co-immunoprecipitation (Co-IP) used 4 µg of Anti-Serine/threonine-protein kinase SNRK2.2/3/4/6/9/10 antibody (PhytoAB, Vestensbergsgreuth, Germany, Cat. No. PHY0715A). For P. Patens samples, 2 µg was used instead. We followed the Invitrogen Dynabeads™ Protein A Immunoprecipitation Kit protocol (Thermo Fischer Scientific) with these exceptions: (1) all steps were performed on ice except for antibody and antigen incubation; (2) elution was performed on a shaker, not on a rotator; (3) the final two washes of step 6 under “Immunoprecipitation of target antigen” used 150 µL 1x PBS for SAG 12.97 samples. For all species except SAG 698-1b, Co-IP was performed seven times per species: 3 biological replicates(cell lysate + SnRK2 antibody); 2 controls which were additionally supplemented with 8M Urea + 1% SDS and 2-4M Urea +0.25-0.5 % SDS, respectively in order to disrupt weak protein-protein or protein-antibody interactions; and 2 bead-only controls (no antibody). For SAG 698-1b, due to poor protein yields, Co-IP was performed on 3 biological replicates, 1 control with 8M Urea + 0.5% SDS, and 1 bead-only control. For P. patens, biological replicates (BRs) corresponded to these ecotypes, respectively: BR1 (Reute), BR2 (Grandsen); and BR3 (Reute); controls mixed both ecotypes. Following the Co-IP, the captured proteins were precipitated according to Wessel and Flügge^103^. In short: protein samples (100 µl) were mixed with 400 µl methanol and vortexed, followed by the addition of 100 µl chloroform and further vortexing. Then, 300 µl water was added, and the mixture was vortexed vigorously for 5 min and centrifuged at 10,000 rpm for 3 min at 4 °C to induce phase separation. The upper aqueous phase was carefully removed, leaving ∼10-20 µl to avoid disturbing the protein interphase. The lower phase was supplemented with 300 µl methanol, vortexed, and centrifuged at 13,000 rpm for 10 min at 4 °C to pellet the protein. The supernatant was discarded, and the protein pellet was air-dried. Protein pellets were resuspended in 40 µl of 0.1% RapiGest SF (Waters Corporation) and vortexed. DTT was added to a final concentration of 5 mM and samples were incubated at 60 °C for 30 min. After adjustment to room temperature, iodoacetamide was added to a final concentration of 15 mM, and samples were incubated in the dark for 30 min. Proteins were digested with trypsin (final concentration 1µg/mL) and incubated at 37 °C for 2.5 hours. Following digestion, samples were acidified with trifluoroacetic acid to ∼0.5% final concentration for adjusting pH to <2. Samples were incubated at 37 °C for 30-45 min to cleave RapiGest. After centrifugation at 13,000 rpm for 10 min, the supernatant was transferred to a new tube and dried in a SpeedVac. Peptides were subsequently desalted (C18 stage tips^104^) and subjected to LC-MS/MS analysis, following the protocol described by Niemeyer et al.^105^

Mass spectrometry raw data files (ThermoScientific .RAW files) were converted to the mzML format using MSConvert and subsequently inspected with MZmine. Since heavy polyethylene glycol (PEG) bleed through was detected^4^, only peptide identification and no quantification was performed. Peptides were identified separately for each file using MetaMorpheus, employing both the GptmTask (Global Post-Translational Modification discovery) and the SearchTask. The maximum number of missed trypsin cleavages was set to 4, while all other parameters were kept at their default settings. For database searches, the following plant protein sequence databases were used: *Arabidopsis thaliana* (TAIR10)^106^, *Closterium peracerosum-strigosum-littorale* complex (NIES68)^107^, *Physcomitrium patens* (v3)^108^, *Mesotaenium endlicherianum* (SAG 12.97 Me1v2 and Chloroplast encoded proteins)^34,88,109^, *Zygnema circumcarinum* (SAG12.97 and UTEX1559 Chloroplast encoded proteins)^72,110^, *Coleochaete scutata* (1KP assembled proteome)^111^. Also, the UniProt cRAP database 2-2-2018 (for common contaminants) and A0A1Y1B8C2, P01870, P54159 (for antibody and bead decontamination) were included. All species-specific protein databases except Arabidopsis (which already includes annotations) were annotated using eggNOG-mapper^112^ using the flags “--sensmode ultra-sensitive −-tax_scope 33090”, see Supplementary Data 3 for results. Peptide scores from the three biological replicates were summed for each protein group. Proteins were categorized based on the presence of peptides across samples: “BR” if peptides were only found in the biological replicate, “L” if also present in SDS-Urea Co-IP data, “A” if found in the no-antibody control.

### SDS-PAGE and Western blot analysis

30 µg of total protein per species was separated on a 10% polyacrylamide gel and transferred onto nitrocellulose membrane using a Mini Trans-Blot® electrophoretic transfer cell (Bio-Rad Laboratories GmbH, Munich, Germany). PageRuler™ Prestained Protein Ladder was used as a molecular weight marker (Thermo Fisher, Cat. No. 26616). Protein transfer was assessed visually with 0.2% ponceau S solution. The membrane was blocked in 1xTBS containing 5% milk powder immediately following ponceau destaining. SnRK2 homologs were detected using anti-SnRK2 PHY0715A (dilution 1:1000) and were visualized by horseradish peroxidase conjugated anti-rabbit secondary antibodies (dilution 1:10000) with a FUSION-SL-4 chemiluminescence imaging system (Vilber Lourmat, Collégien, France) in supersaturation mode for faint band visualization.

### Phylogenetics

For all phylogenies, the protein databases used for the Co-IP were used. For ABF, sequences that were found using a blastp search with e-value cut-off of 0.1 with as query “Me1_v2_0220640.1” were used for tree construction. For PP2C, RAF, and SnRK2 analysis, we built on the expanded repertorie of sequenced streptophyte genomes^113^ and protein databases of the following organisms were additionally added and combined to one database^114^: *Zea mays*, *Pica abies*, *Saleginella moellendroffi, Azolla filiculoides, Anthoceros agrestis*, *Marchanthia polymorpha*, *Spirogloea muscicola*, *Chara braunii*, *Mesostrima viride*, *Klebsormium nites*, *Chlorokybus atmophycus*, *Coleochaete scutate*, *Chlamydomonas reinhardtii, Volvox carteri, Ulva mutabilis*. For PP2C, all proteins that were identified with HMMER search with the motif “PF00481.24” and an e-value cut-off of 0.1 were used. For RAF, sequences that were found using a blastp search with e-value cut-off of 10^−50^ with as query “Me1_v2_0218750” were used for tree construction. For SnRK, sequences that were found using a blastp search with percent identify cut-off of 50% and alignment length of minimal 250 AA with as query “Me1_v2_0141810.1” were used initially. Due to poorly resolved tree structure, the sequence of Zea, Marchanthia, Volvox and Spirogloea were removed, and the following sequences were added: the SnRK2s from Amborella trichpoda, Ostreococcus lucimarinus, Penium margaritaceum and the following SnRK2s obtained from transcriptome shotgun assembly database from NCBI: JV767425:SnRK2_Nitella_mirabilis, GICF01002155:Spirogyra_pratensis, GJZN01009725:Mougeotiopsis_calospora, GBSL01050164.1:Coleochaete_orbicularis, JO159324.1:Chaetosphaeridium_globosum. For all 4 phylogenetic analysis, the selected protein sequences were aligned using MAFFT with the L-INS-i algorithm and a maximum likelihood phylogenetic tree was constructed using IQ-Tree with 1,000 ultrafast bootstrap replicates. The following models were used: SnRK2 & RAF = JTTDCMut+F+G4, ABF= VT+F+R5, PP2C= LG+I+G. The resulting phylogenies were visualised in iTOL^115^.

### Software used

blast 2.16.0 (Johnson et al.^116^) Bowtie2 (Langmead and Salzberg^117^), ChimeraX (Pettersen et al.^118^), eggNOG-mapper v2.1.8 (Huerta-Cepas et al.^112,119^), FUNC-E 2.0.1 (https://github.com/SystemsGenetics/FUNC-E), HMMER 3.3.2 (http://hmmer.org/), IGV 2.14.0 (Robinson et al.^120^), iqtree 2.1.3 (Minh et al.^121^), iTOL v7 (Letunic and Bork^115^), imputeLCMD (Cosmin Lazar (2015). imputeLCMD: A collection of methods for left-censored missing data imputation. R package version 2.0. http://CRAN.R-project.org/package=imputeLCMD) integrated into Perseus), MACS2 (Zhang et al.^122^), MAFFT^123^ 7.304b, MaxQuant^124^ v1.6.3.4, MetaMorpheus^125^ CMD 1.0.0, MSconvert^126^ v3, mzmine^127^ v4.3.0, Perseus^128^ v1.6.14.0, Python 3.9.7 (packages used: argsparse, copy, numpy, math, matplotlib, pandas, scipy.stats, seaborn, statistics; https://docs.python.org/release/3.9.7/), Thermo Xcalibur 4.8, and R^129^ v4.3.2 (<u>Packages used:</u> htmlwidgets, ggpmisc, ggplot2, ggrepel, Homer^130^, plotly, readr, shiny, shinylive).

## DATA AVAILABILITY

The mass spectrometry proteomics data have been deposited to the ProteomeXchange Consortium via the PRIDE^131^ under Project accession PXD078633 (Co-IP) and PXD078585 (phosphoproteomic data).

All supplementary data have been deposited on Zenodo under doi: 10.5281/zenodo.20328447

## ACKNOWLEDGEMENTS

J.d.V. thanks (a) the European Research Council for funding under the European Union’s Horizon 2020 research and innovation program (Grant Agreement No. 852725; ERC-StG “TerreStriAL”) as well as the Horizon Europe programme (Grant Agreement No. 101230161; ERC-CoG “StreptoProgram”) and (b) the German Research Foundation (DFG) for support through project “SHOAL” (514060973), project “Klebsome” (509535047) and the framework of the Priority Program “MAdLand – Molecular Adaptation to Land: Plant Evolution to Change” (SPP 2237; 440231723 and 528076711), in which C.F.K. and J.M.S.Z. partake as associate members. J.M.S.Z. and C.F.K. are grateful for support through the IMPRS Genome Science. We extend our gratitude to Dr. Tatyana Darienko for growing *Zygnema* and *Coleochaete*. We are very greatful for the help of Merle Aden and Isabell Peymann for their support with the western blot. We are very grateful for the help of Dr. Maaike Bierenbrootspot with script optimisation. The Q Exactive/Ultimate 3000 LC-MS system was granted by DFG INST 186/1230-1 FUGG and for the hormone measurements I.F. acknowledges funding by DFG INST 186/822-1. K.S. is financed by DFG INST 186/1465-1. H.N. was supported by the Max Planck Society.

## CONTRIBUTIONS

J.Z., G.R., A.H., S.C.S., L.H., H.N. and J.d.V. were involved in either experimental planning, excecution, and/or data analysis of the phosphoproteomics experiments. J.Z., M.S., J.C.M., S.A. and J.d.V. were involved in the apocarotanoid measurements. J.Z., A.Z., I.M.R., J.S., J.d.V., were involved in either experimental planning, excecution, and/or data analysis of the DNA Affinity Purification Sequencing. J.Z., K.S., C.D., O.V. and J.d.V. were involved in the experimental excecution, the measurements and/or data analysis of the co-immunoprecipitaiton experiment. J.Z., S.K., C.F.K., I.F., and J.d.V. were involved in the plant hormone measurements. J.Z. performed the phylogenetic analysis. All authors discussed the results and contributed to the final manuscript. J.Z., M.S., A.M., and J.d.V. were involved in protein-protein interaction studies.

