## Supplementary Fig. 1 for "Homologous ABA-independent kinase tracks coalesced into osmotic stress circuits during plant terrestrialization"

**Supplementary Figure** 1.

Data distribution and correlation of phosphopeptide intensities.

**A.** Histogram showing the distribution of the complete dataset. Observed values are shown in blue, while imputed values are displayed in other colors. The number associated with each imputation group indicates the number of missing values in the corresponding condition and corresponding phosphopeptide. Normality was assessed for the observed data combined with imputed datasets 1–4 using scipy.stats.normaltest.

**B.** Correlation matrices for both datasets.

**C.** Histograms showing the data distribution for each individual sample.

*A*

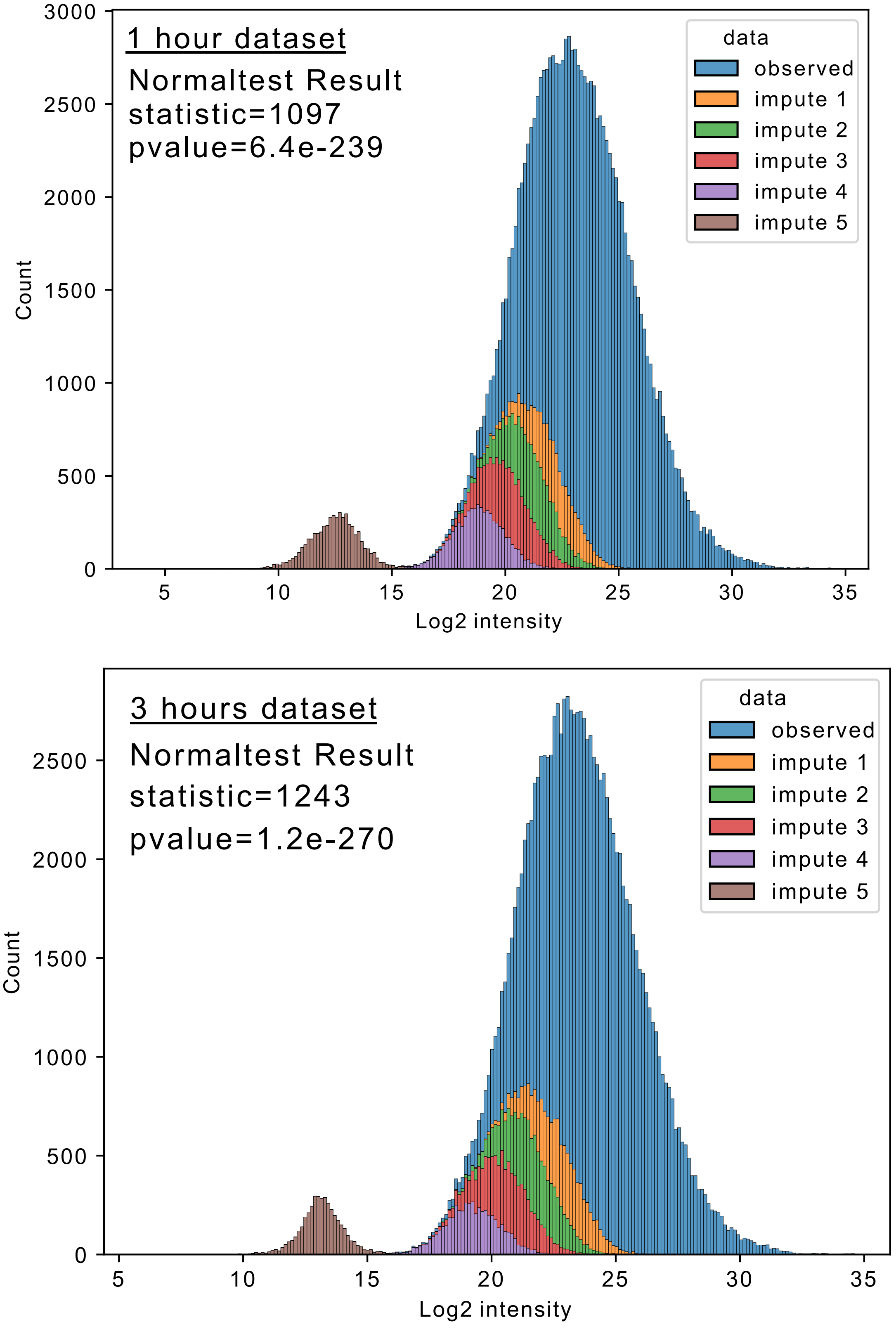

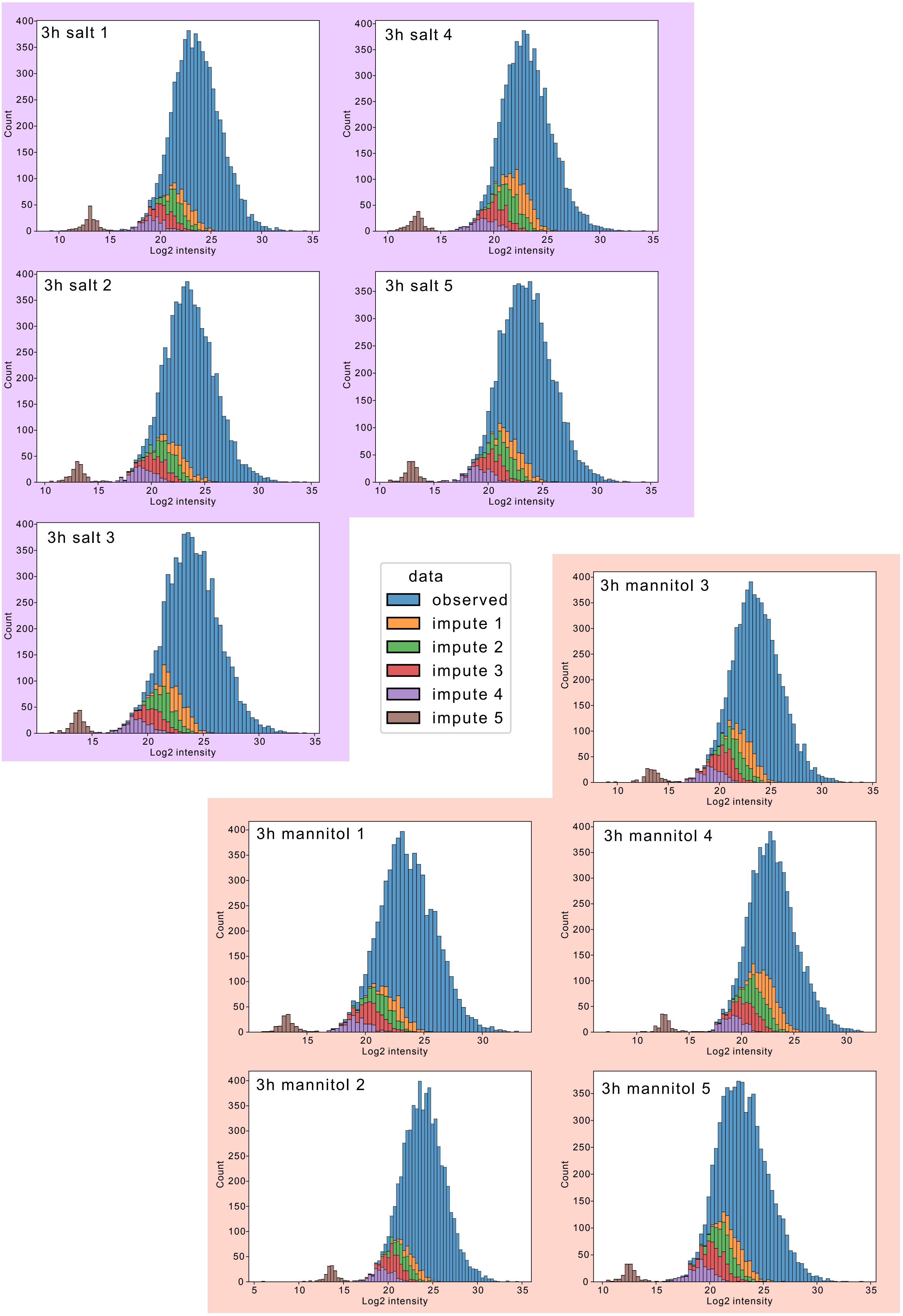
3h salt 1
3h salt 2
3h salt 3
3h salt 4
3h salt 5
3h mannitol 1
3h mannitol 2
3h mannitol 3
3h mannitol 4
3h mannitol 5
Count
Log2 intensity
data
observed
impute 1
impute 2
impute 3
impute 4
impute 5

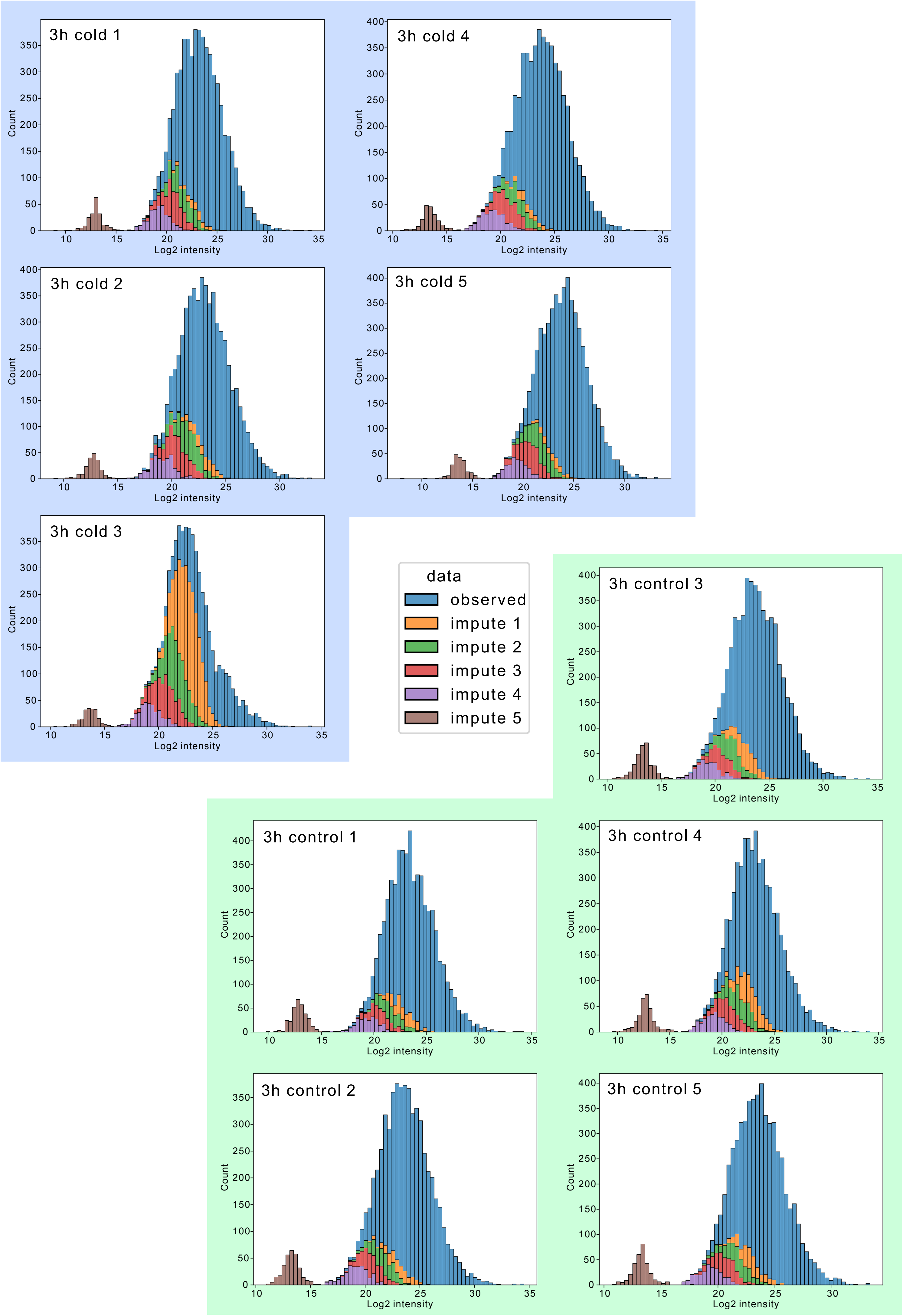
3h cold 1
3h cold 2
3h cold 3
3h cold 4
3h cold 5
3h control 1
3h control 2
3h control 3
3h control 4
3h control 5
Count
Log2 intensity
data
observed
impute 1
impute 2
impute 3
impute 4
impute 5

**B**

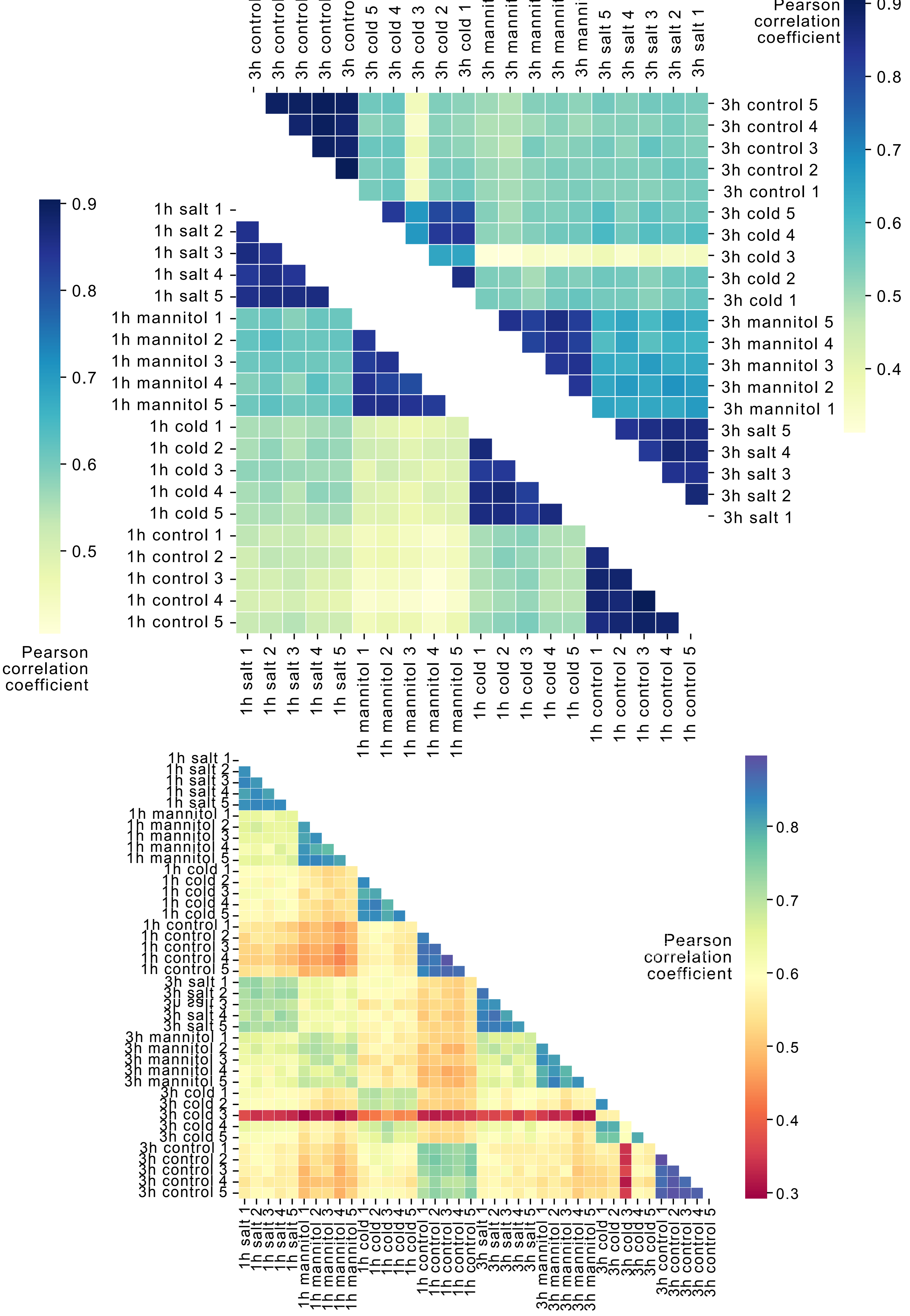
Pearson correlation coefficient
0.9
0.8
0.7
0.6
0.5
0.4
0.3
1h salt 1
1h salt 2
1h salt 3
1h salt 4
1h salt 5
1h mannitol 1
1h mannitol 2
1h mannitol 3
1h mannitol 4
1h mannitol 5
1h cold 1
1h cold 2
1h cold 3
1h cold 4
1h cold 5
1h control 1
1h control 2
1h control 3
1h control 4
1h control 5
3h salt 1
3h salt 2
3h salt 3
3h salt 4
3h salt 5
3h mannitol 1
3h mannitol 2
3h mannitol 3
3h mannitol 4
3h mannitol 5
3h cold 1
3h cold 2
3h cold 3
3h cold 4
3h cold 5
3h control 1
3h control 2
3h control 3
3h control 4
3h control 5

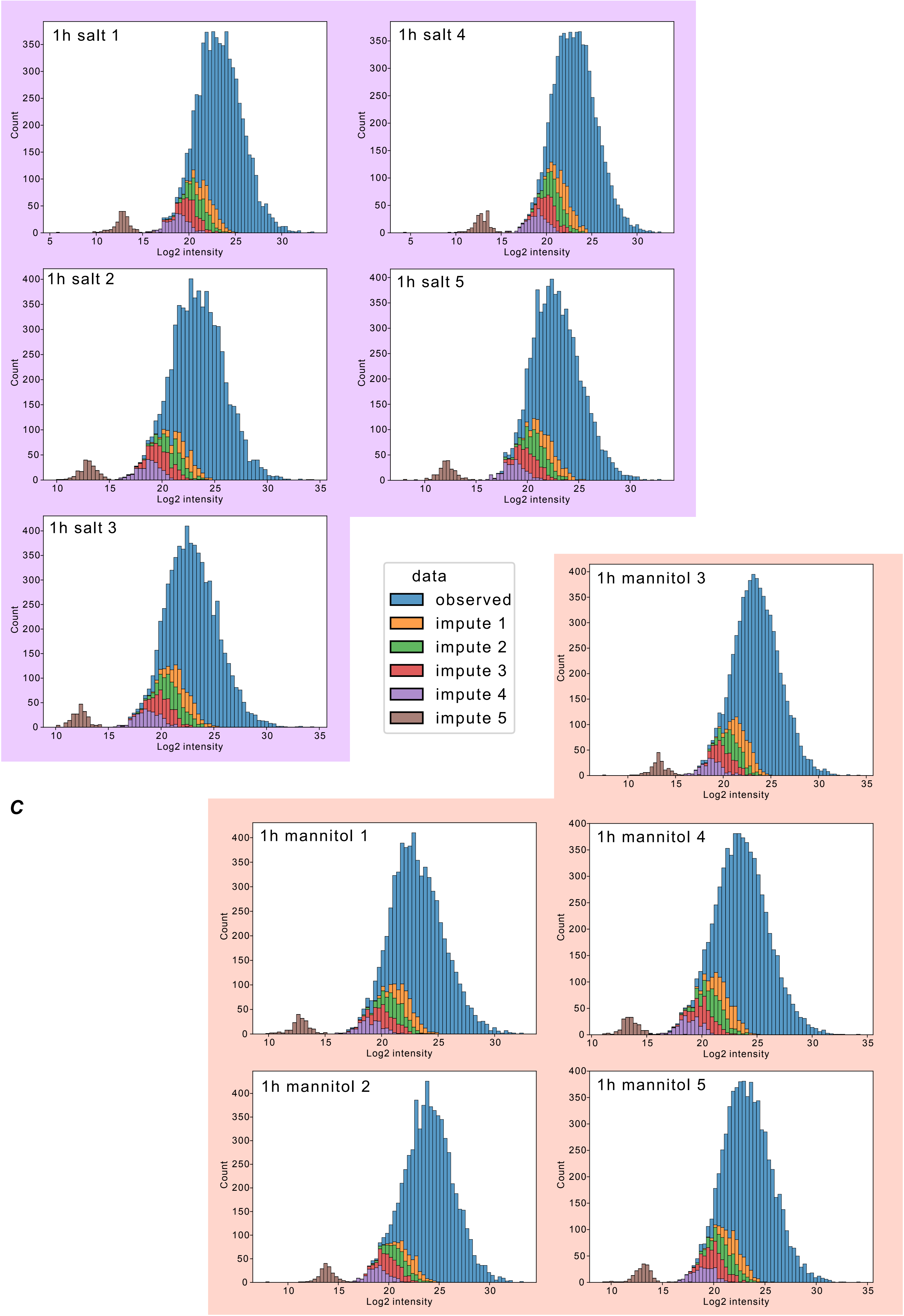

1h salt 1
1h salt 4
1h salt 2
1h salt 5
1h salt 3
Count
Log2 intensity
data
observed
impute 1
impute 2
impute 3
impute 4
impute 5
C
1h mannitol 3
1h mannitol 1
1h mannitol 4
1h mannitol 2
1h mannitol 5

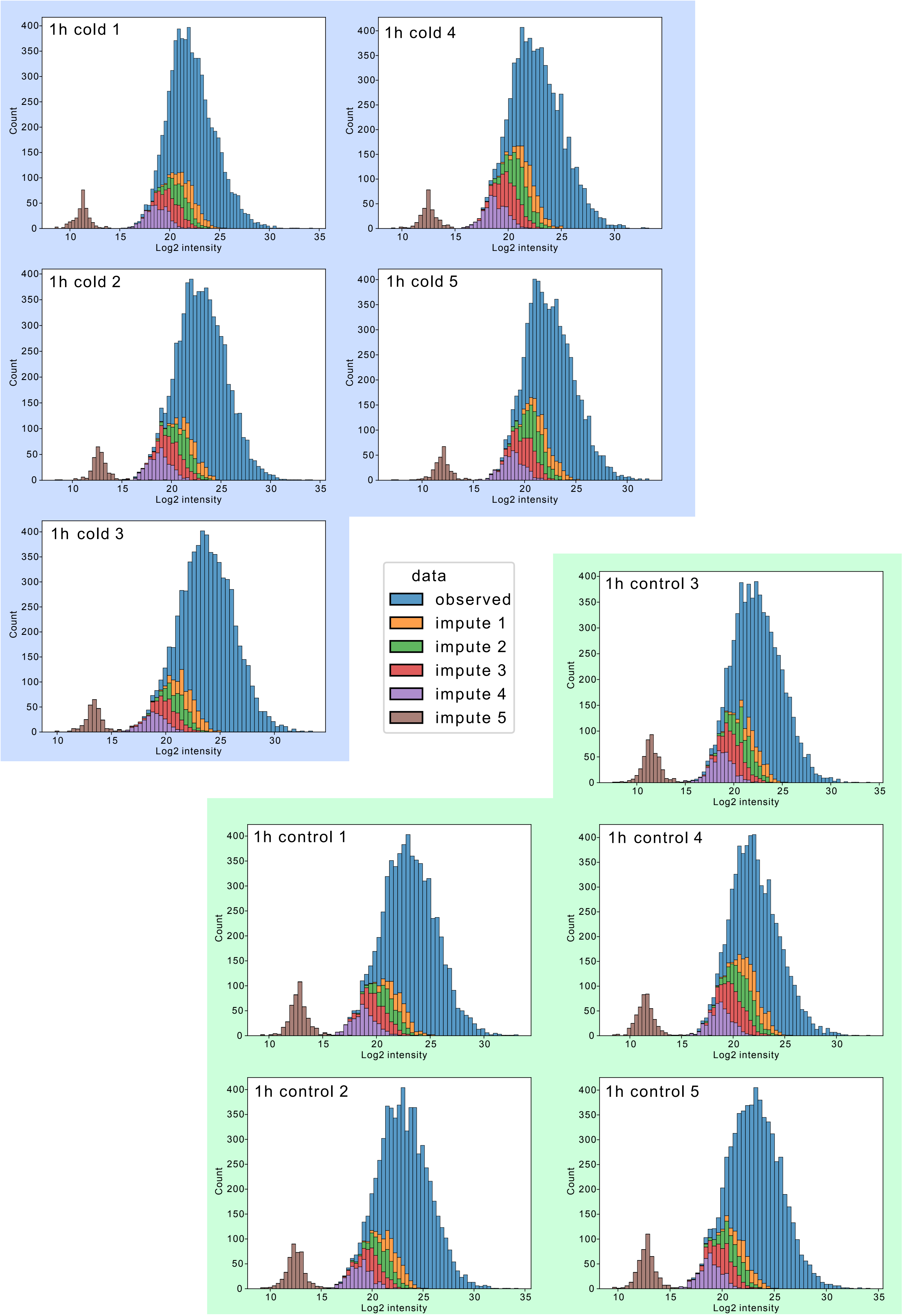
1h cold 1
1h cold 2
1h cold 3
1h cold 4
1h cold 5
1h control 1
1h control 2
1h control 3
1h control 4
1h control 5
Count
Log2 intensity
data
observed
impute 1
impute 2
impute 3
impute 4
impute 5
