## Supplementary Fig. 2 for "Homologous ABA-independent kinase tracks coalesced into osmotic stress circuits during plant terrestrialization"

Supplementary Figure 2.
Phylogenetic Analysis of Histidine Kinases in the green lineage

**Annotation**
Coloured ranges are annotated with the use the histidine kinases of Arabidopsis thaliana that have all been annotated and the OsHK (Osmotic histidine kinase) from Mesotaenium that has been identified in this study. Protein domains are annotated using hmmscan (HMMER 3.3.2) and the Pfam database. Transmembrane (TM) regions were found with TMHMM 2.0.

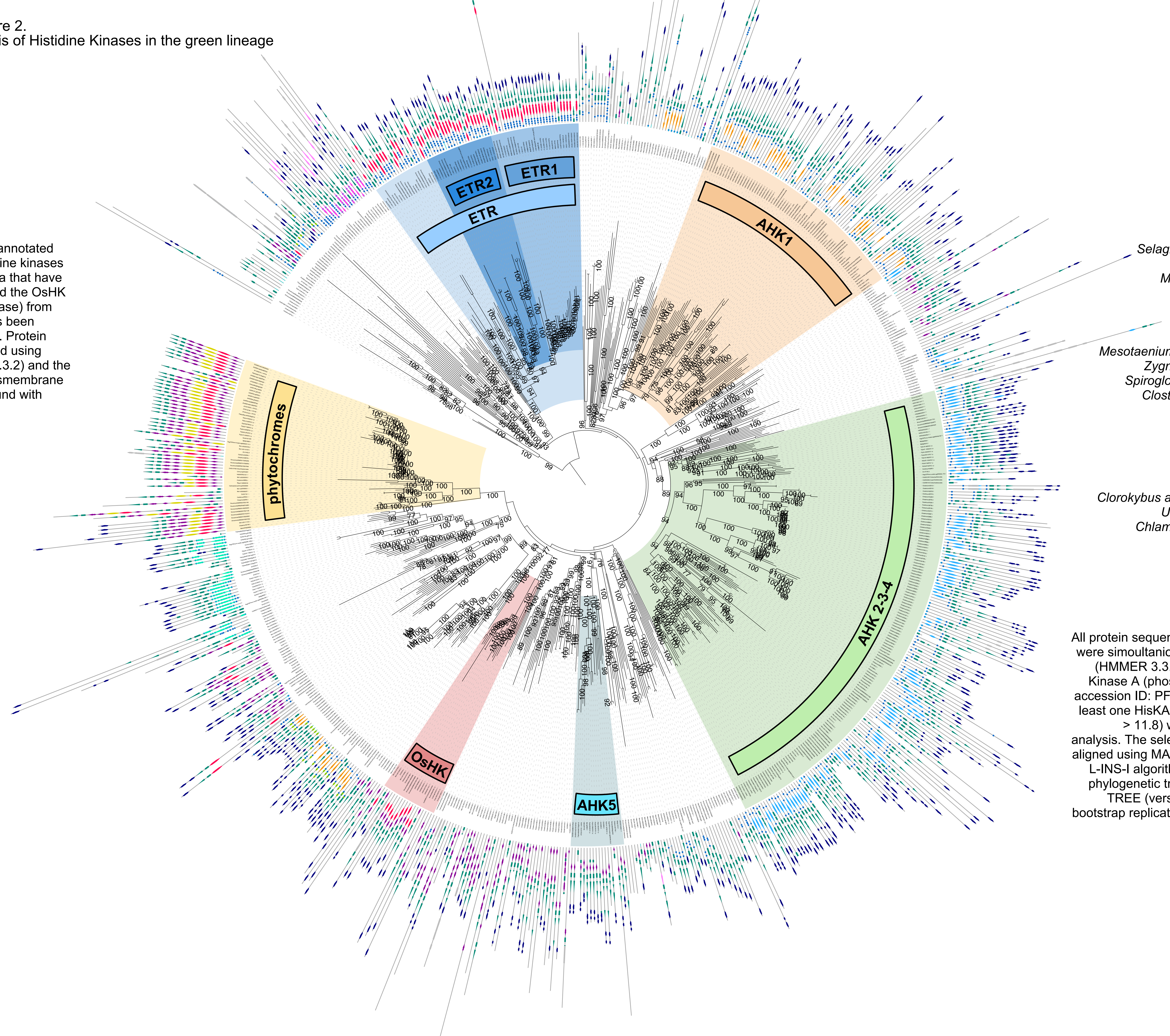


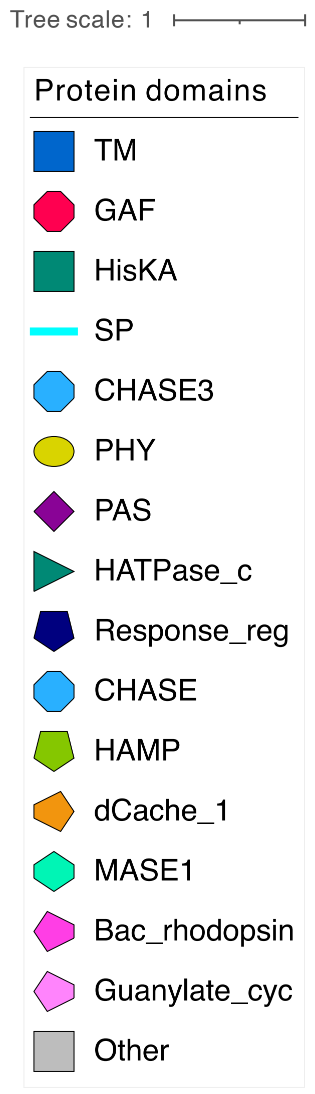


**Species**

Embryophates:
*Arabidopsis thaliana* ID = AT
*Zea mays* ID = Zm
*Pica Abies* ID = Pabies
*Selaginella moellendorfii* ID = Smoel
*Azolla filiculoides* ID = Azfi
*Marcanthia polymorpha* ID = Mp
*Physcomitrium patens* ID = Pp
*Anthoceros agrestis* ID = Aagr

Streptophyte algae:
*Mesotaenium endlicherianum* ID = Me1_v2
*Zygnema circumcarinatum* ID = Zci
*Spirogloea muscicola* ID = Smuscicola
*Closterium* NIES68 ID = Closterium
*Chara braunii* ID = Cbraunii
*Coleochaete scutata* ID = Cs
*Klebsormidium nitens* ID = Kfl
*Mesostigma viride* ID = Mesvi

Chlorophytes:
*Clorokybus atmophycus* ID = Catmophycus
*Usnea mutabilis* ID = Umutabilis
*Chlamydomonas reinhardtii* ID = Cre
*Volvox carteri* ID = Vocar

**Methods**
All protein sequences from all selected species were simoultaniouly scanned with hmmsearch (HMMER 3.3.2) for the presence of the His Kinase A (phospho-acceptor) domain (Pfam accession ID: PF00512.31). Only those with at least one HisKA domain (E-value < 6.3, score > 11.8) were kept for the pyhlogenetic analysis. The selected protein sequences were aligned using MAFFT (version 7.304b) with the L-INS-I algorithm and a maximum likelihood phylogenetic tree was constructed using IQ-TREE (version 2.1.3) with 1,000 ultrafast bootstrap replicates and ModelFinder selecting the Q.pfam+R10 model.
