## Supplementary Fig. 3 for "Homologous ABA-independent kinase tracks coalesced into osmotic stress circuits during plant terrestrialization"

### Supplementary Figure(s) 3. DAPseq with ABF

**Figure 3a.** All signals (generated with MACS2, OneDup) and predicted summits (generated with MACS2, autoDup) are visualized on the Mesotaenium genome, using gene annotations from version 2. Regions ±60 kb around peaks with scores > 200 are shown. Peaks adjacent to mitochondrial genes or located within the last 10 kb of a contig are excluded. Gene expression patterns and gene names are displayed for those with a $\log_2$ fold change (L2FC) > 1 and a maximum TPM > 1.5.

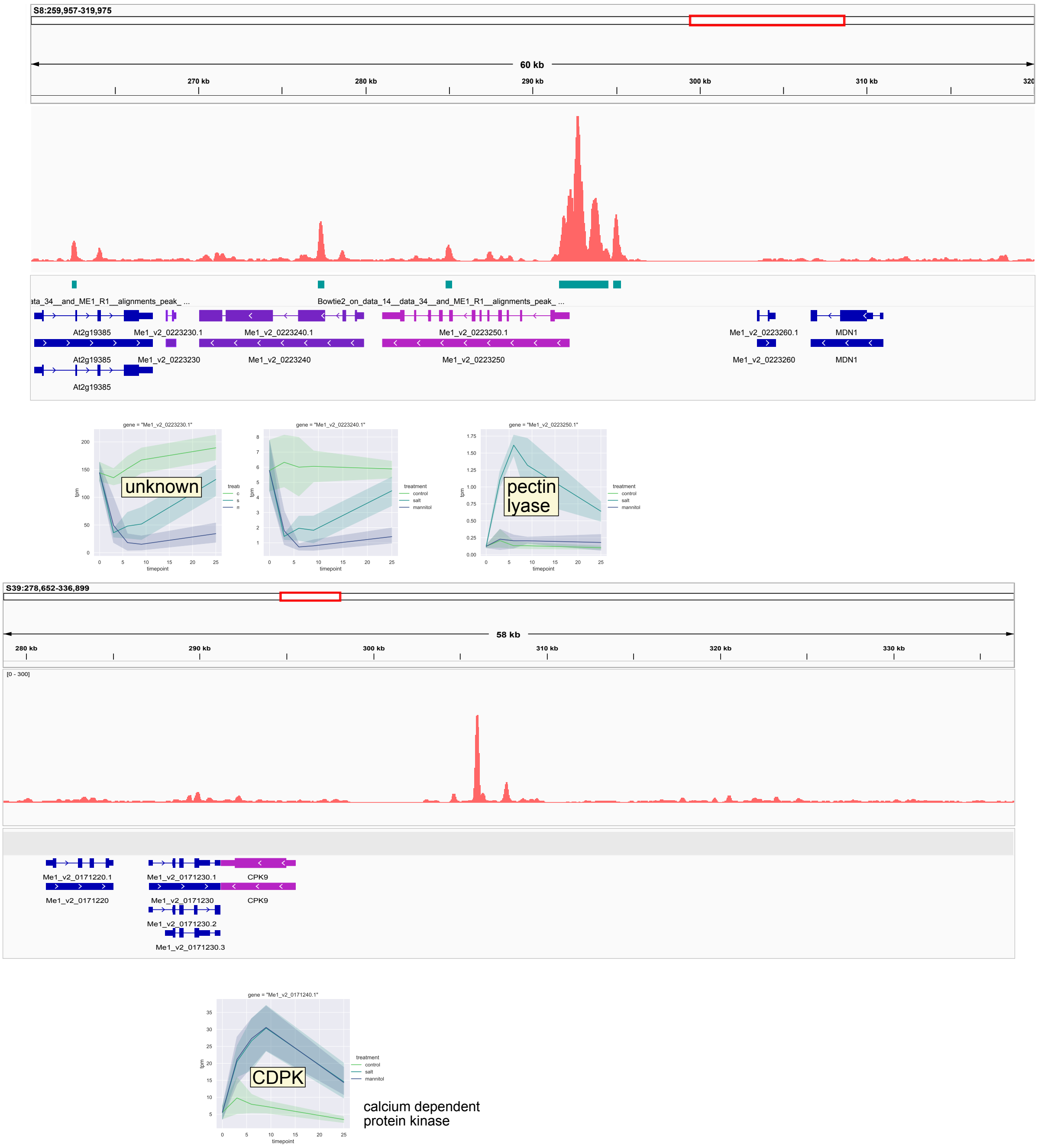

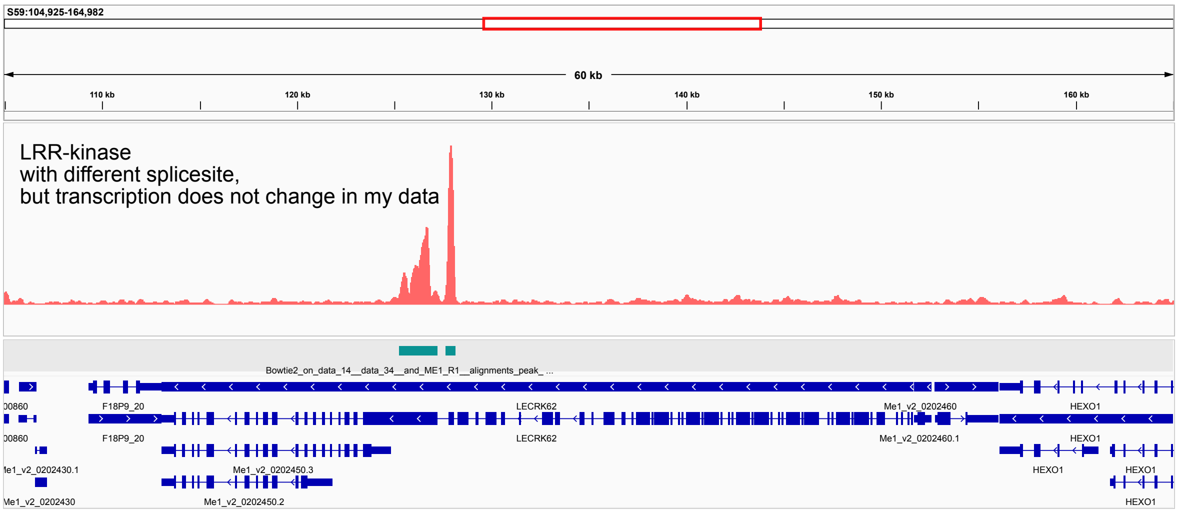
S59:104,925-164,982
60 kb
110 kb
120 kb
130 kb
140 kb
150 kb
160 kb
LRR-kinase
with different splicesite,
but transcription does not change in my data
Bowtie2_on_data_14__data_34__and_ME1_R1__alignments_peak_...
F18P9_20
LECRK62
Me1_v2_0202460
HEXO1
Me1_v2_0202460.1
Me1_v2_0202450.3
Me1_v2_0202450.2
Me1_v2_0202430

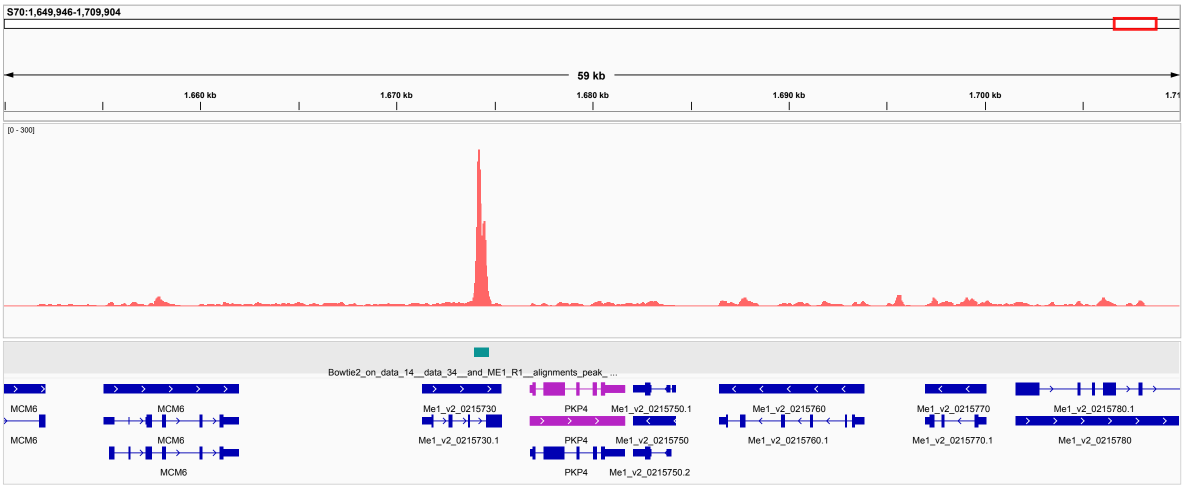
S70:1,649,946-1,709,904
59 kb
1.660 kb
1.670 kb
1.680 kb
1.690 kb
1.700 kb
[0 - 300]
Bowtie2_on_data_14__data_34__and_ME1_R1__alignments_peak_...
MCM6
Me1_v2_0215730
Me1_v2_0215730.1
PKP4
Me1_v2_0215750.1
Me1_v2_0215750
Me1_v2_0215750.2
Me1_v2_0215760
Me1_v2_0215760.1
Me1_v2_0215770
Me1_v2_0215770.1
Me1_v2_0215780.1
Me1_v2_0215780

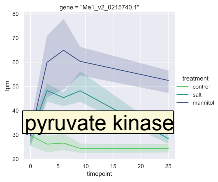
pyruvate kinase
treatment
control
salt
mannitol
timepoint

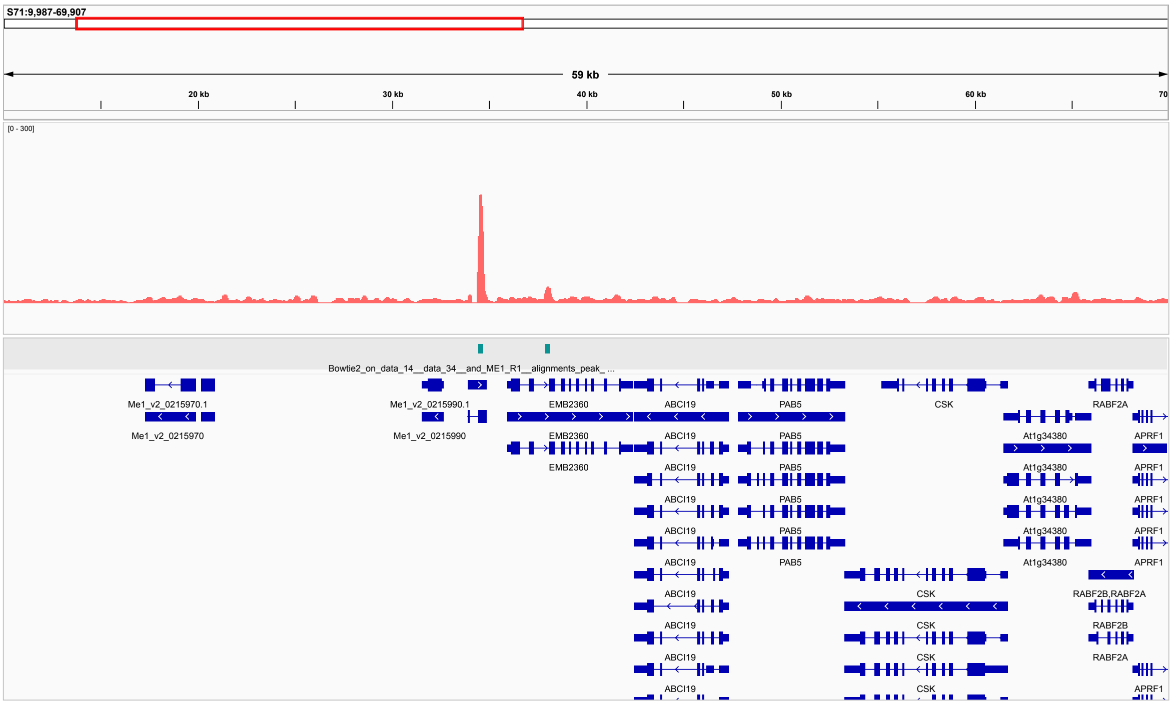
S71:9,987-69,907
59 kb
20 kb
30 kb
40 kb
50 kb
60 kb
[0 - 300]
Bowtie2_on_data_14__data_34__and_ME1_R1__alignments_peak_...
Me1_v2_0215970.1
Me1_v2_0215970
Me1_v2_0215990.1
Me1_v2_0215990
EMB2360
ABCI19
PAB5
CSK
RABF2A
At1g34380
APRF1
RABF2B,RABF2A
RABF2B

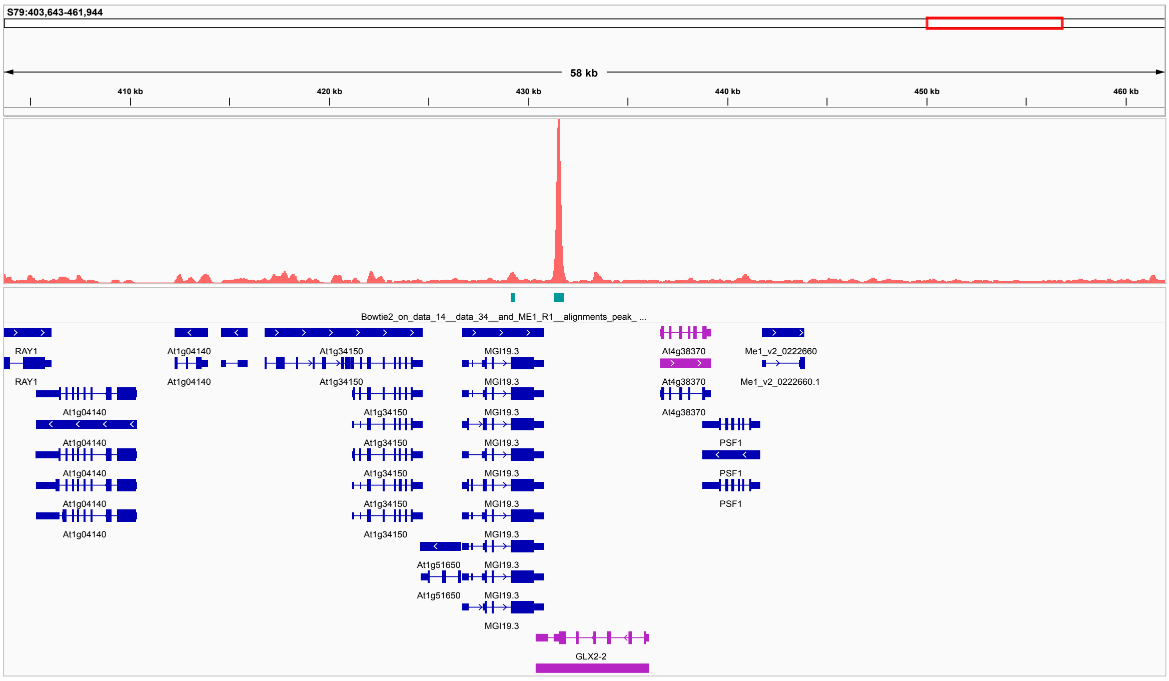


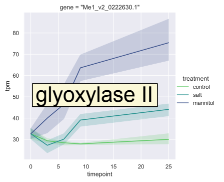


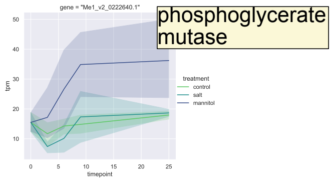


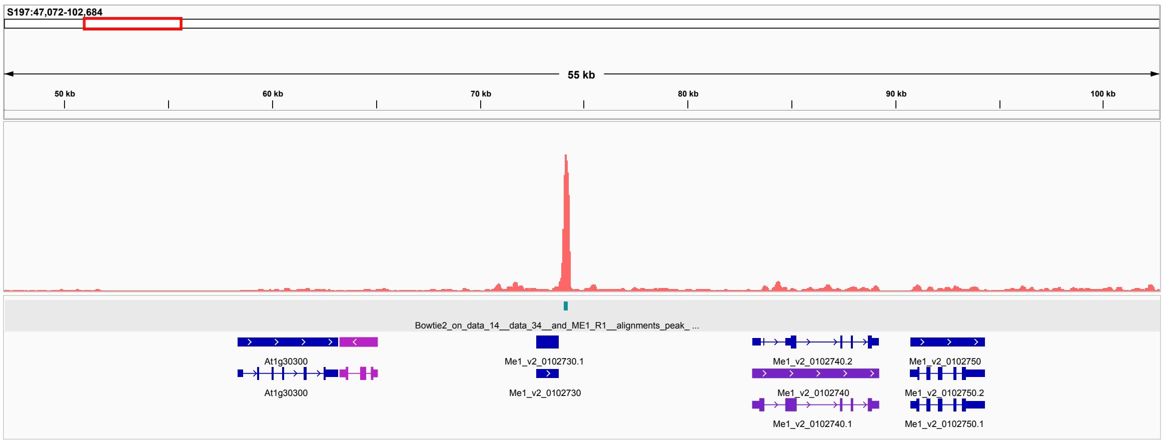


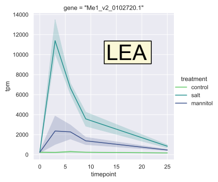


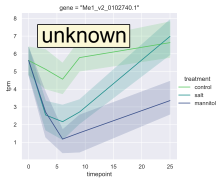


LEA/ABA-inducible protein PHV A1-like, unknown

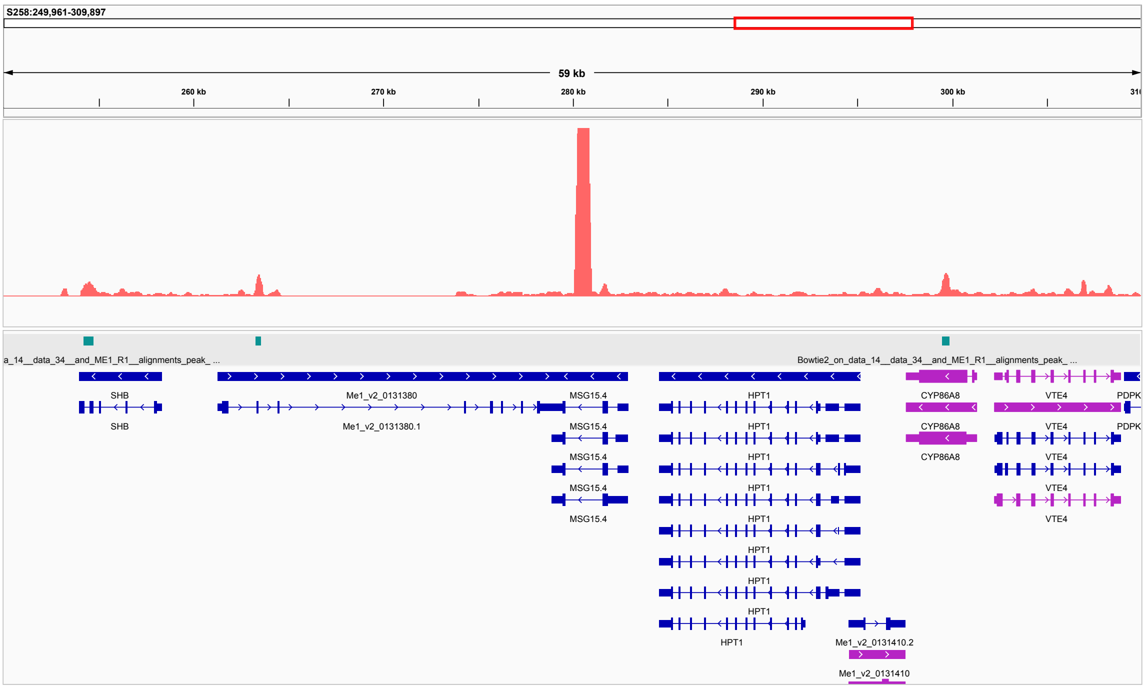


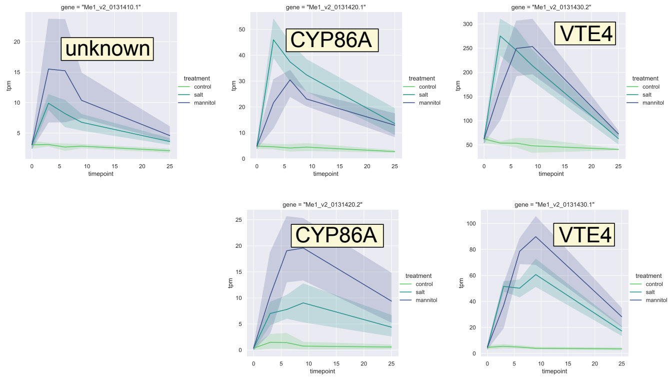


cutin biosynthesis (in At)

vitamin E biosynthesis

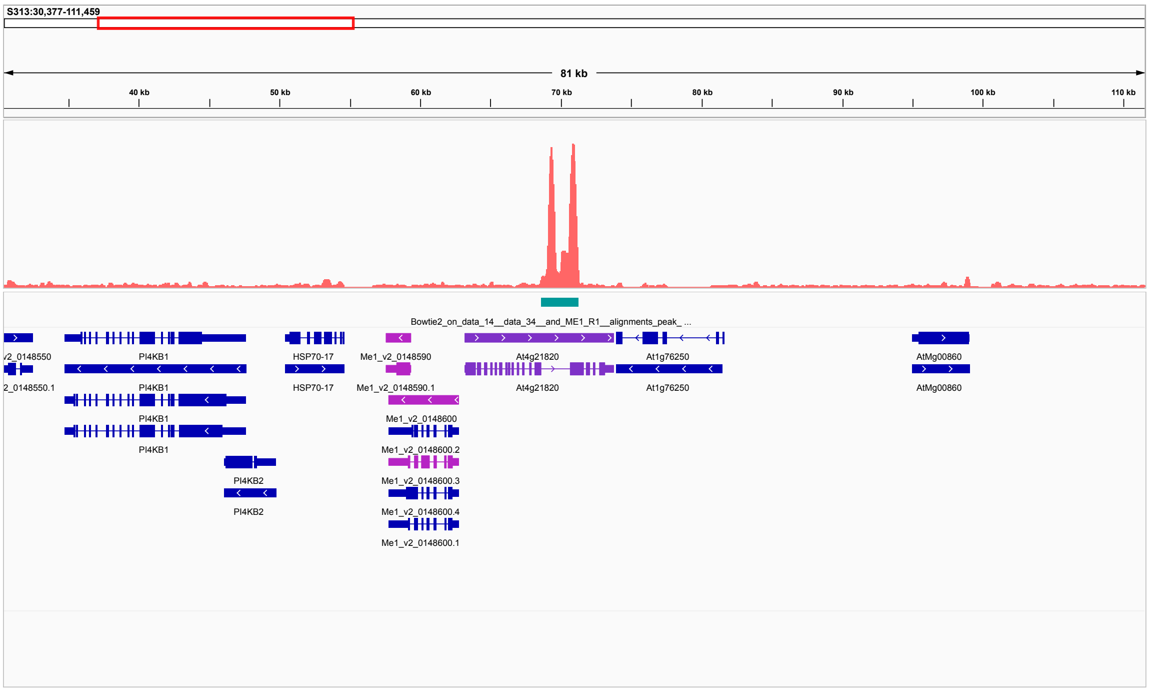


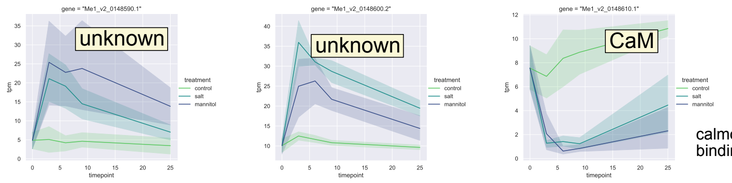


calmoduling
binding protein: Ca signaling

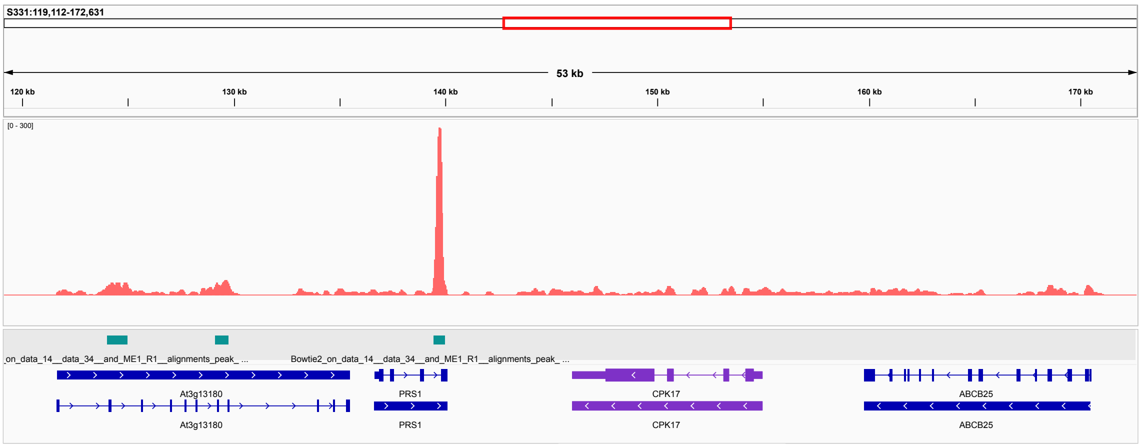


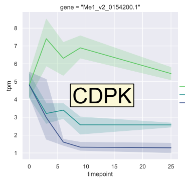


calcium dependent protein kinase

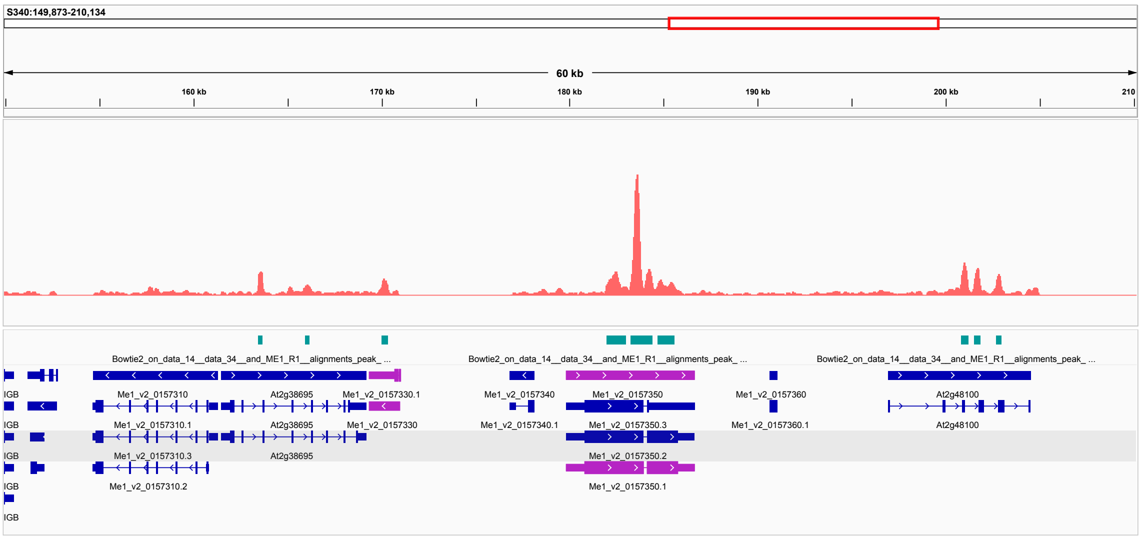


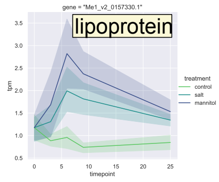


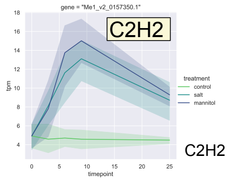


C2H2 zinc finger

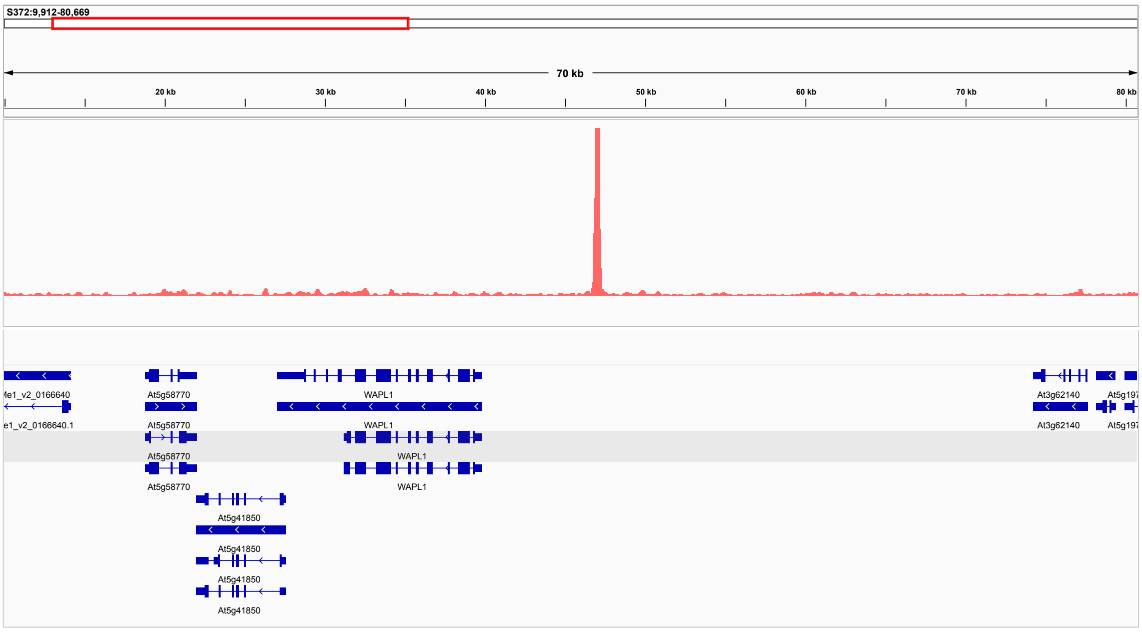

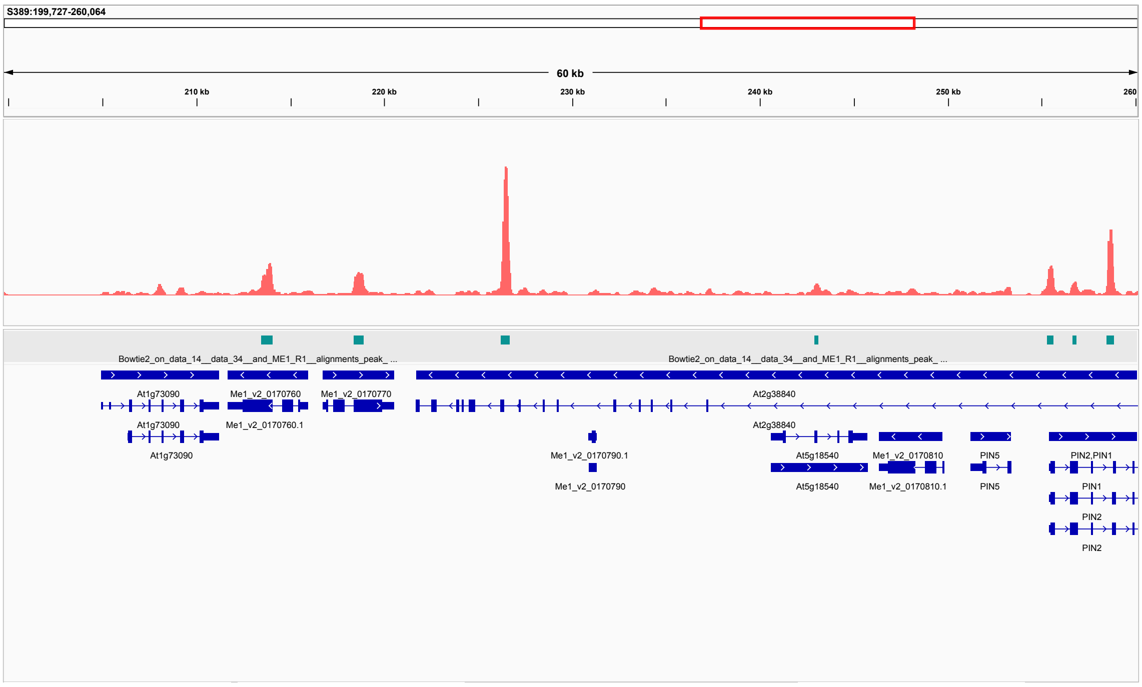


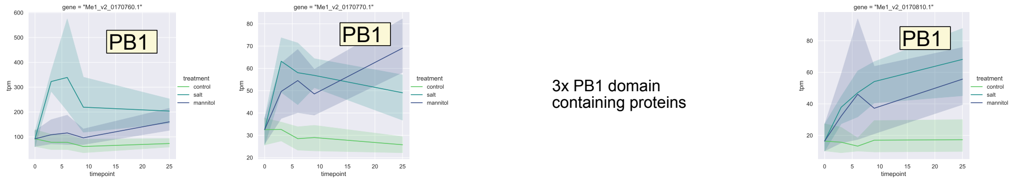


3x PB1 domain containing proteins

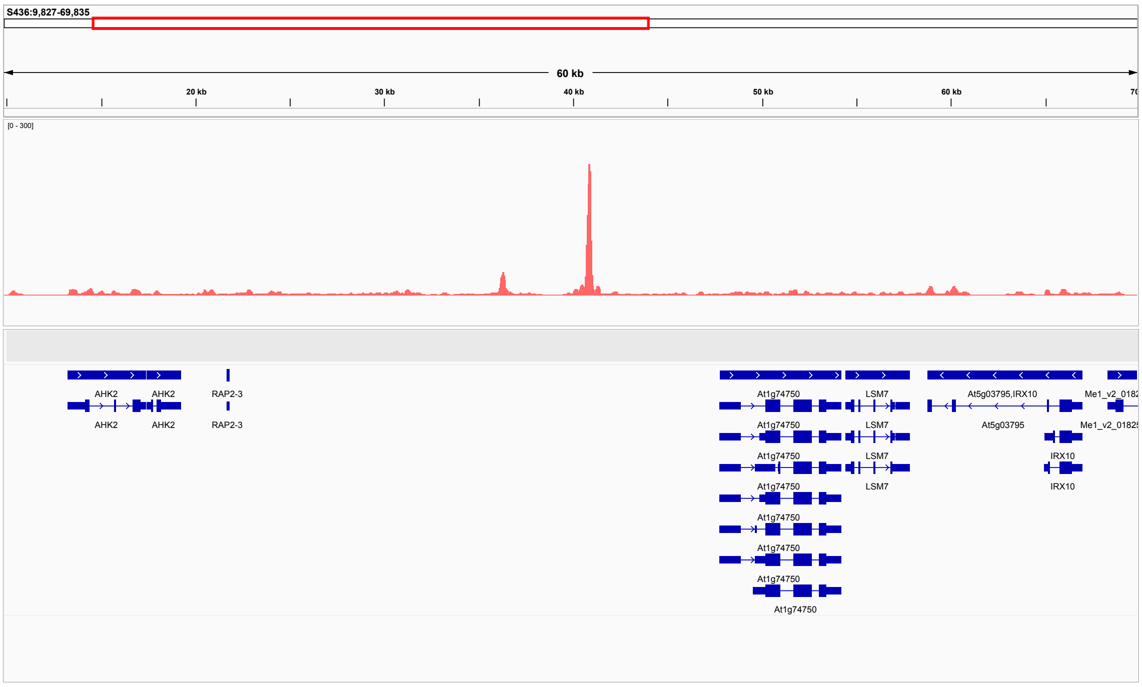


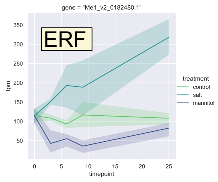


ethylene-responsive transcription factor

**Figure 3b.** Comparison of peaks located upstream of genes with gene expression under osmotic stress. Peaks were predicted using OneDup.

Top panel: treatment with 0.8 M mannitol; bottom panel: treatment with 0.15 M NaCl.

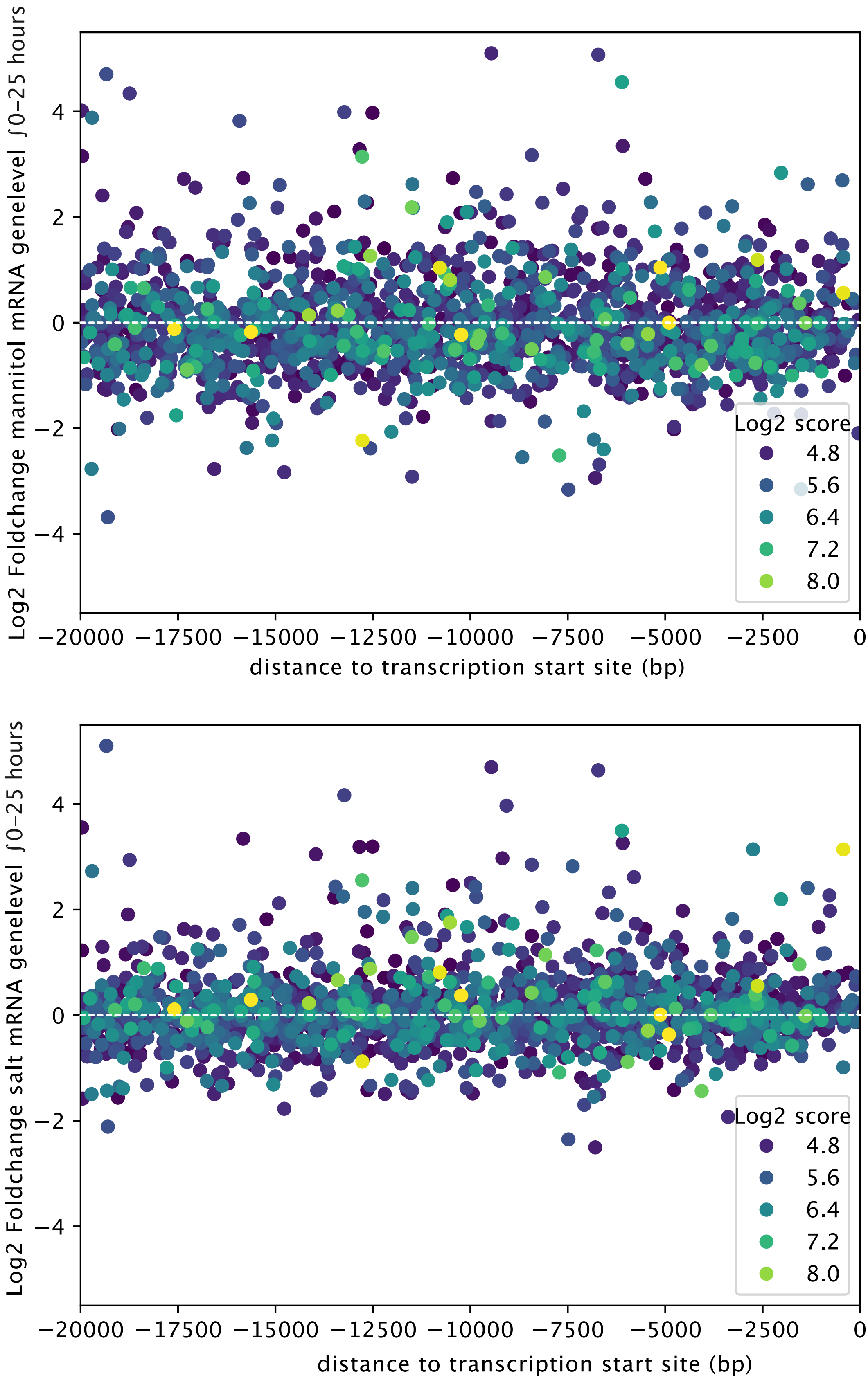
