## Supplementary Fig. 4 for "Homologous ABA-independent kinase tracks coalesced into osmotic stress circuits during plant terrestrialization"

**B-G)** Distribution of total protein levels and comparisons of total protein across conditions. Extended figure discription on other pages.

***B***

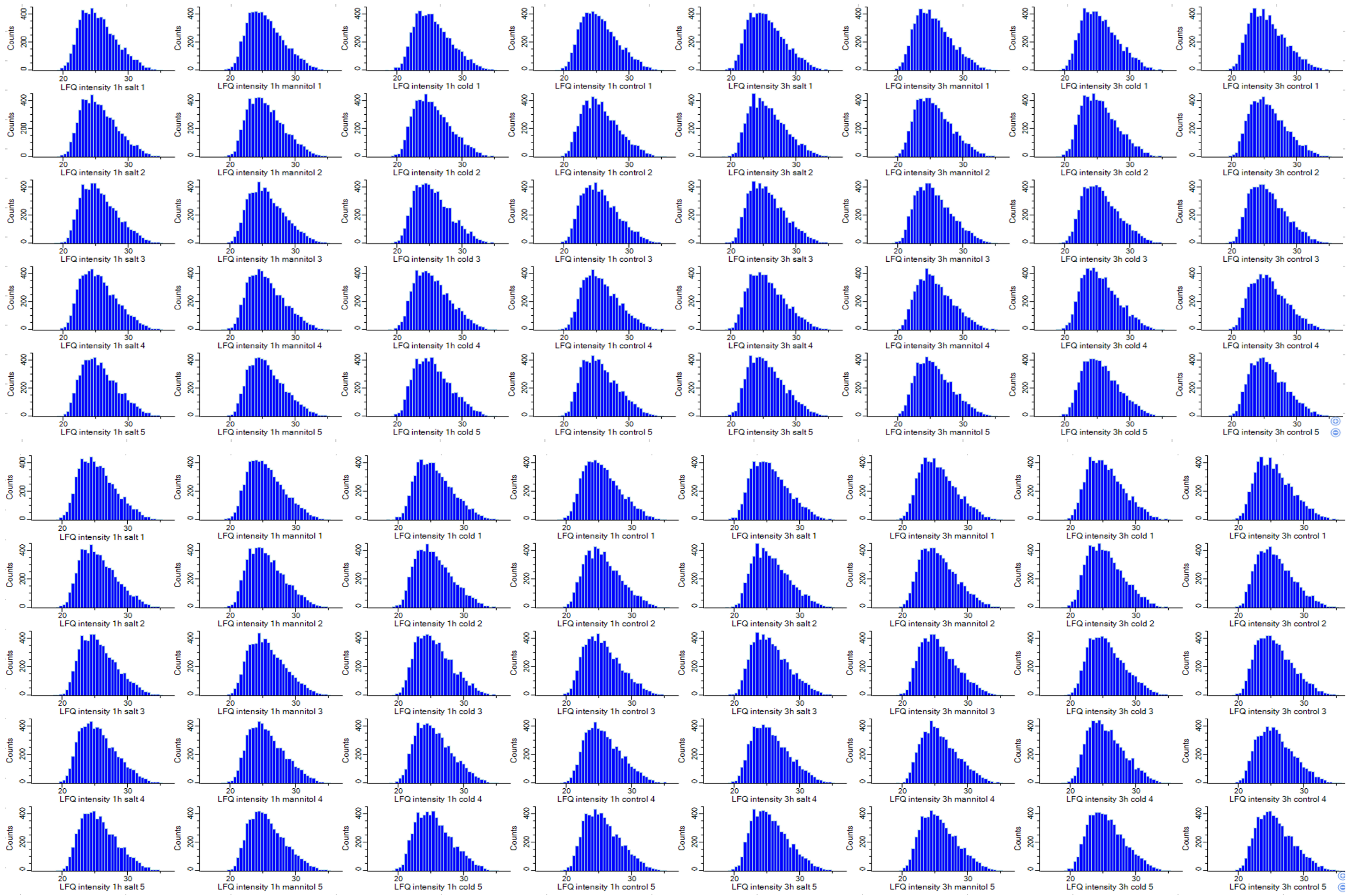


***C***

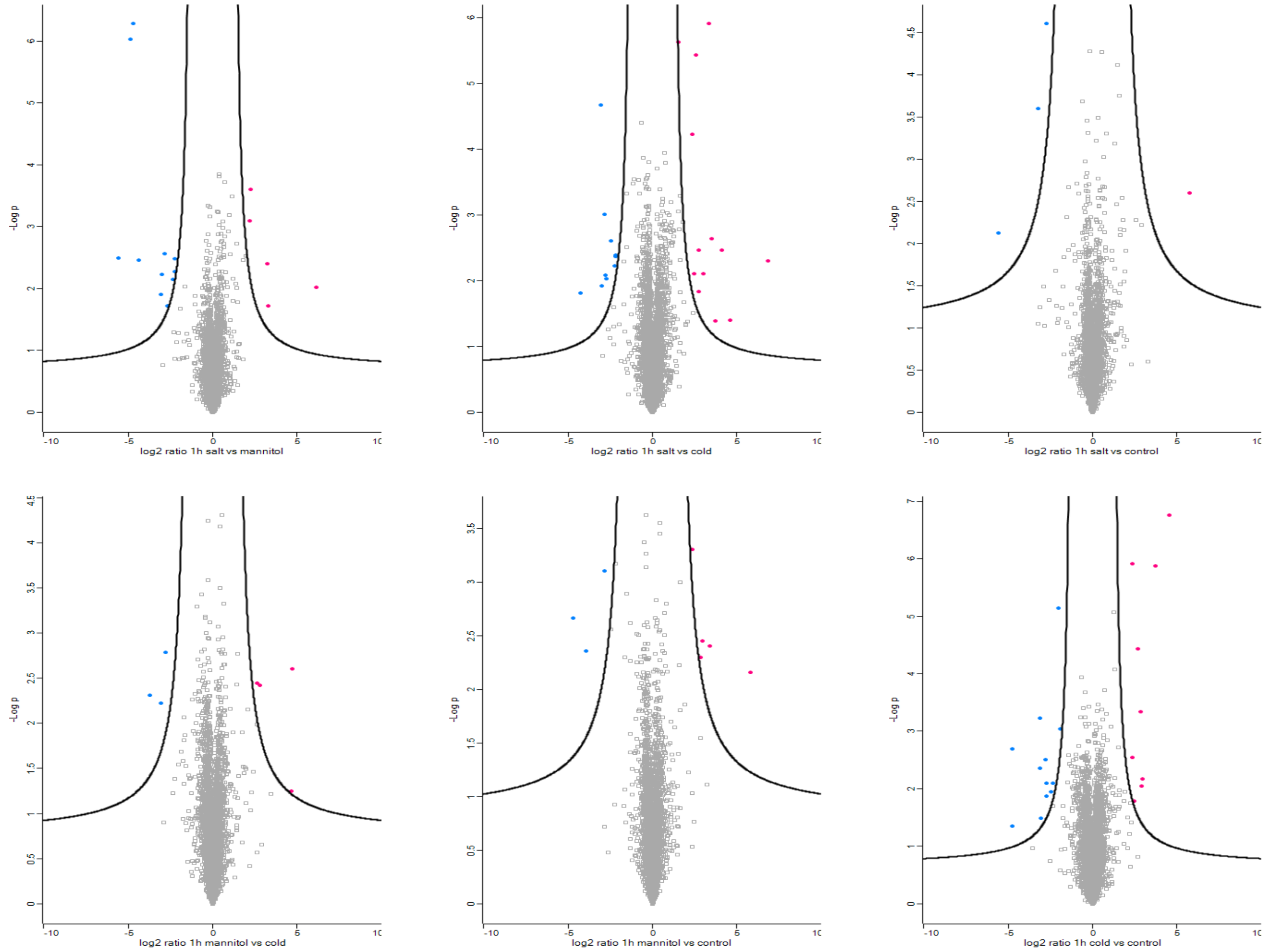

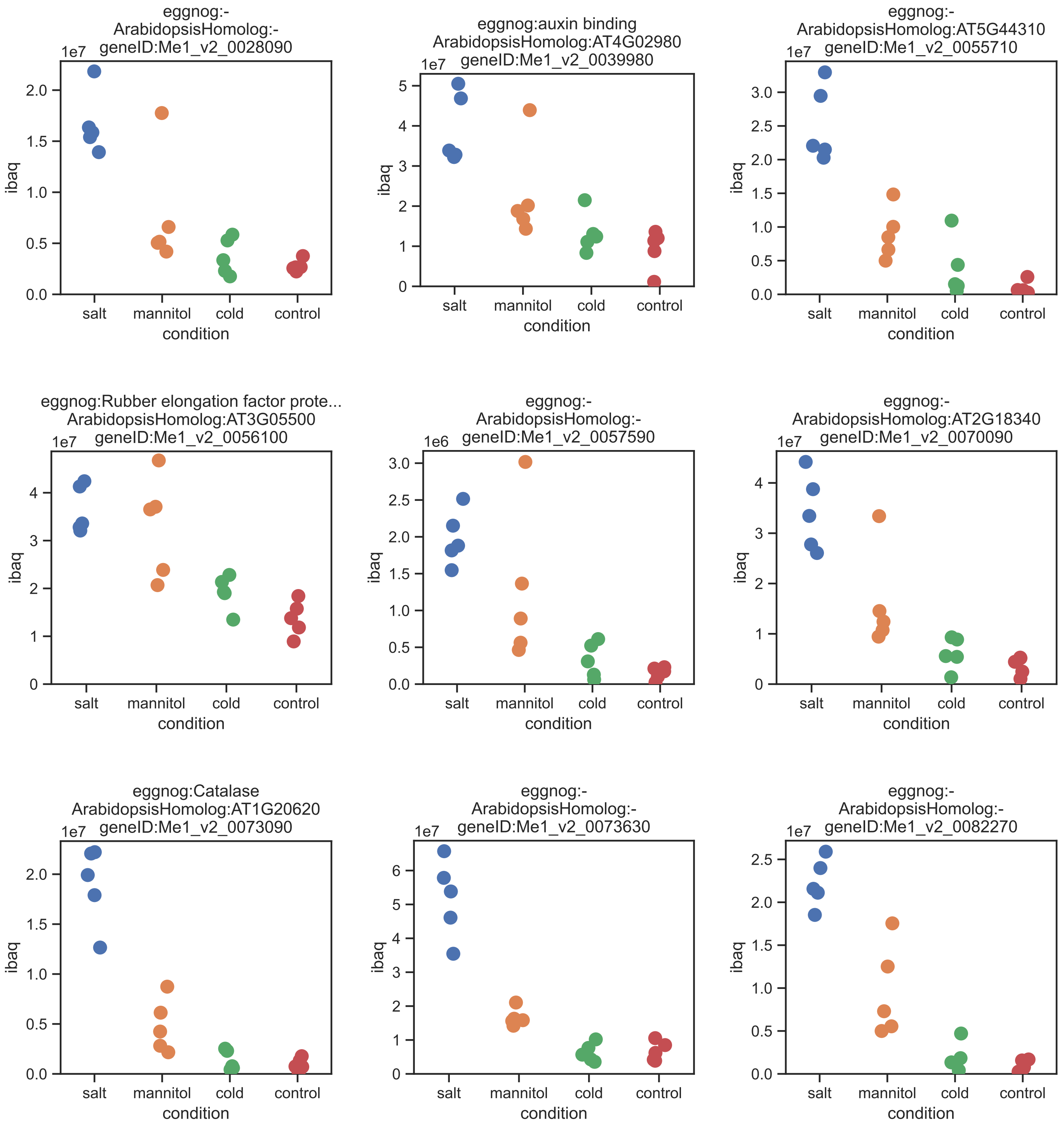

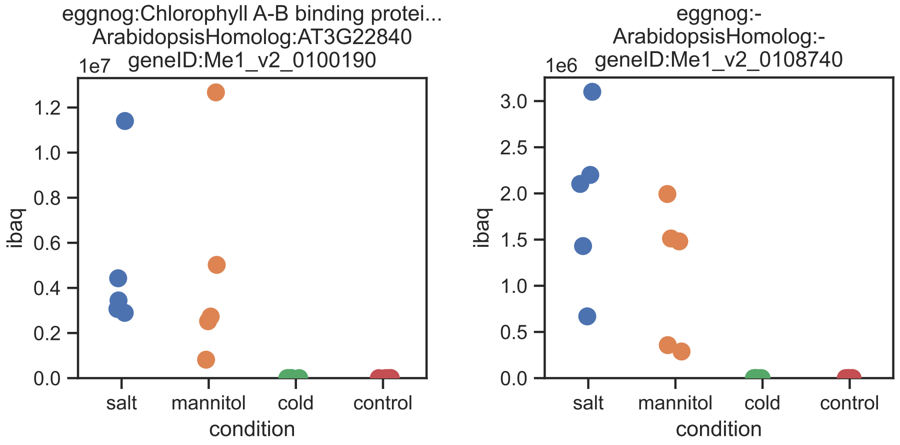


**F+G.** iBAQ values (without imputation) for all proteins showing strong significant changes, as determined by ANOVA-based pairwise comparisons of LFQ values with Benjamini–Hochberg–adjusted p-values (FDR < 0.01). The corresponding gene ID of *Mesotaenium's* protein, its closest *Arabidopsis thaliana* homolog (if any), and the automatic eggNOG annotation are given.

**F** = 1 hour experiment.
**G** = 3 hours experiment.

***D***

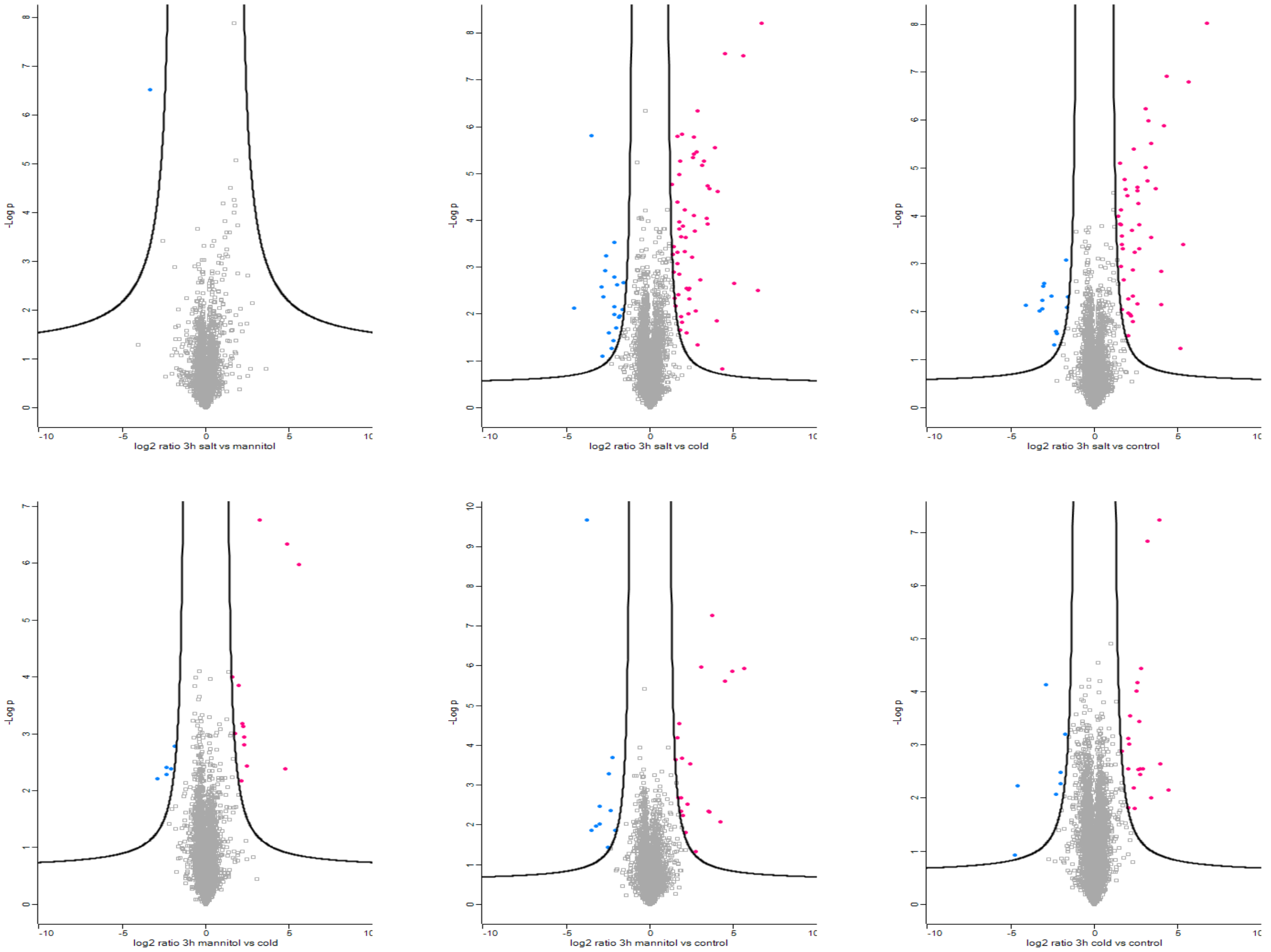


**B.** Distribution plot of log2 LFQ values per sample.

**C-E.** Volcano plots: pairwise comparisons of LFQ samples between conditions. Significantly (t-test with FDR < 0.05) different proteins are coloured.

**C** = comparisons different conditions at 1 hour.

**D** = comparisons different conditions at 3 hours.

**E** = comparisons 1 versus 3 hours within the same condition.

***E***

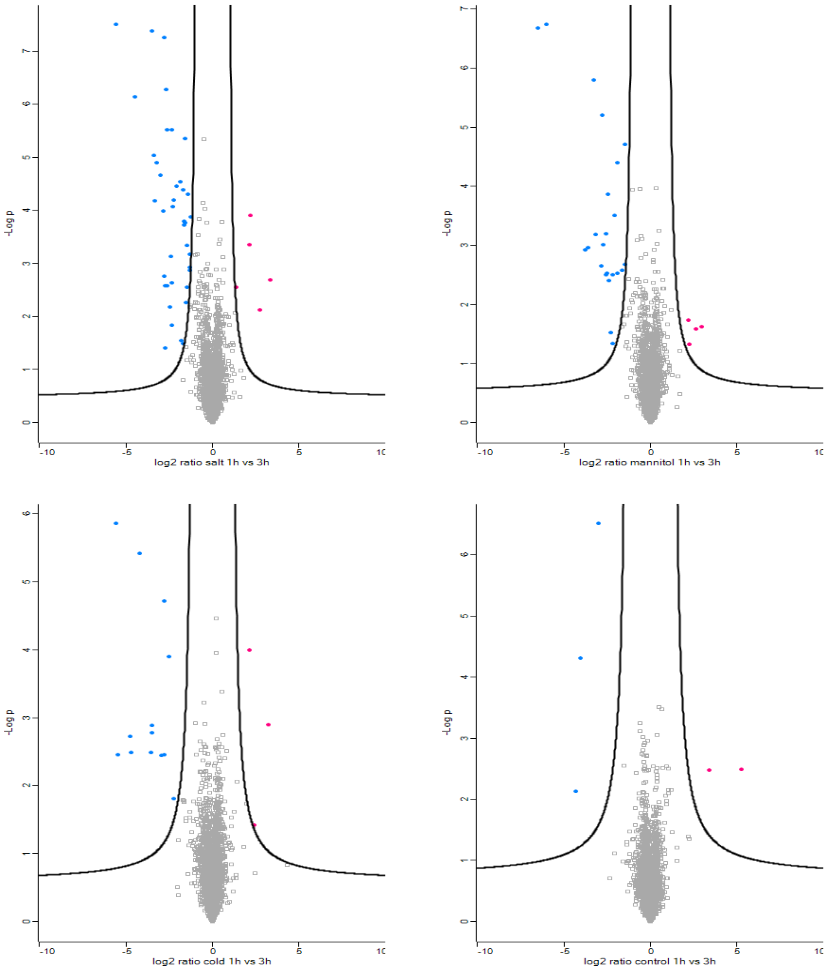

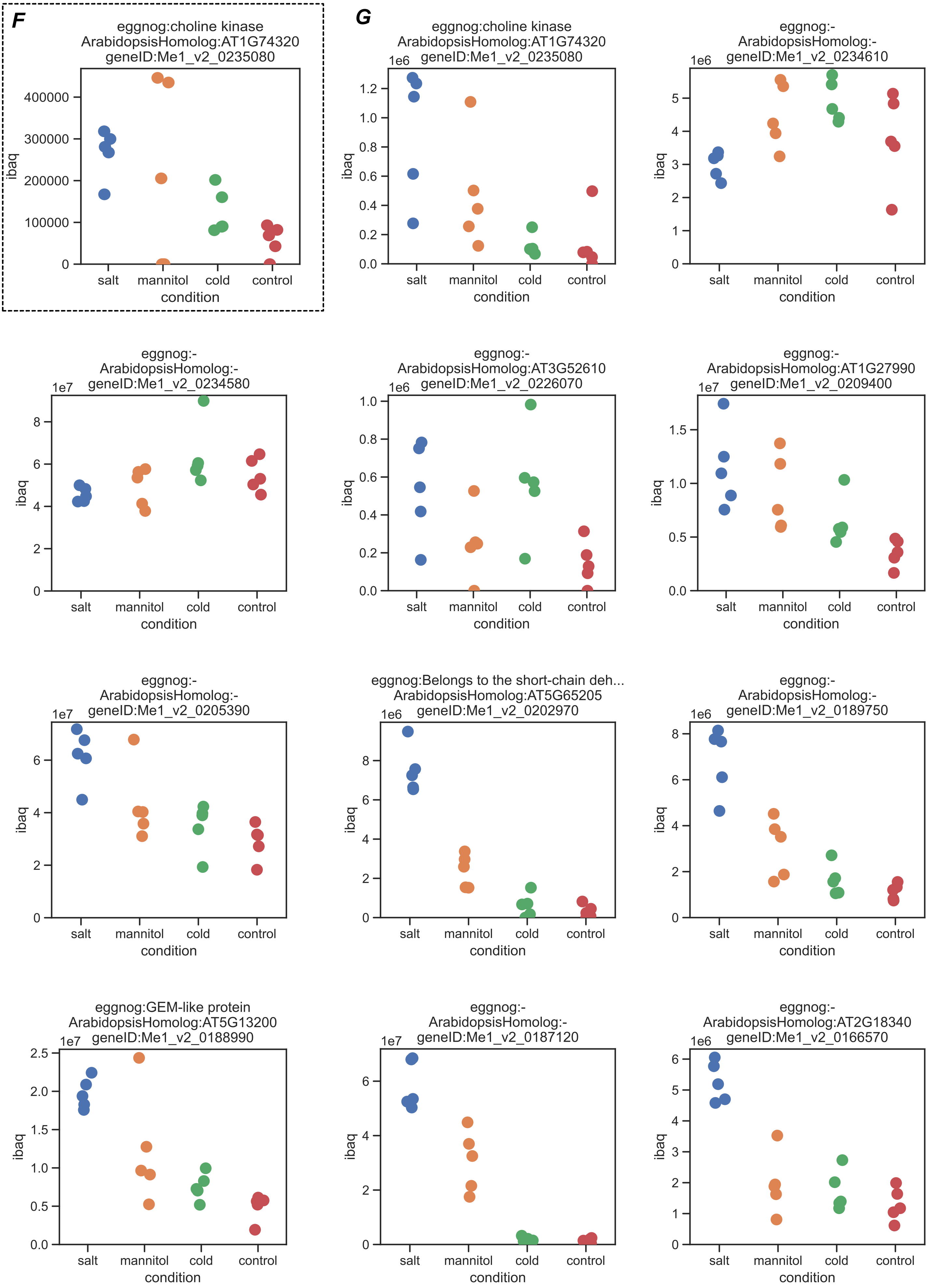

F
eggnog:choline kinase
ArabidopsisHomolog:AT1G74320
geneID:Me1_v2_0235080
ibaq
salt
mannitol
cold
control
condition
G
eggnog:choline kinase
ArabidopsisHomolog:AT1G74320
geneID:Me1_v2_0235080
1e6
eggnog:-
ArabidopsisHomolog:-
geneID:Me1_v2_0234610
1e6
eggnog:-
ArabidopsisHomolog:-
geneID:Me1_v2_0234580
1e7
eggnog:-
ArabidopsisHomolog:AT3G52610
geneID:Me1_v2_0226070
1e6
eggnog:-
ArabidopsisHomolog:AT1G27990
geneID:Me1_v2_0209400
1e7
eggnog:-
ArabidopsisHomolog:-
geneID:Me1_v2_0205390
1e7
eggnog:Belongs to the short-chain deh...
ArabidopsisHomolog:AT5G65205
geneID:Me1_v2_0202970
1e6
eggnog:-
ArabidopsisHomolog:-
geneID:Me1_v2_0189750
1e6
eggnog:GEM-like protein
ArabidopsisHomolog:AT5G13200
geneID:Me1_v2_0188990
1e7
eggnog:-
ArabidopsisHomolog:-
geneID:Me1_v2_0187120
1e7
eggnog:-
ArabidopsisHomolog:AT2G18340
geneID:Me1_v2_0166570
1e6

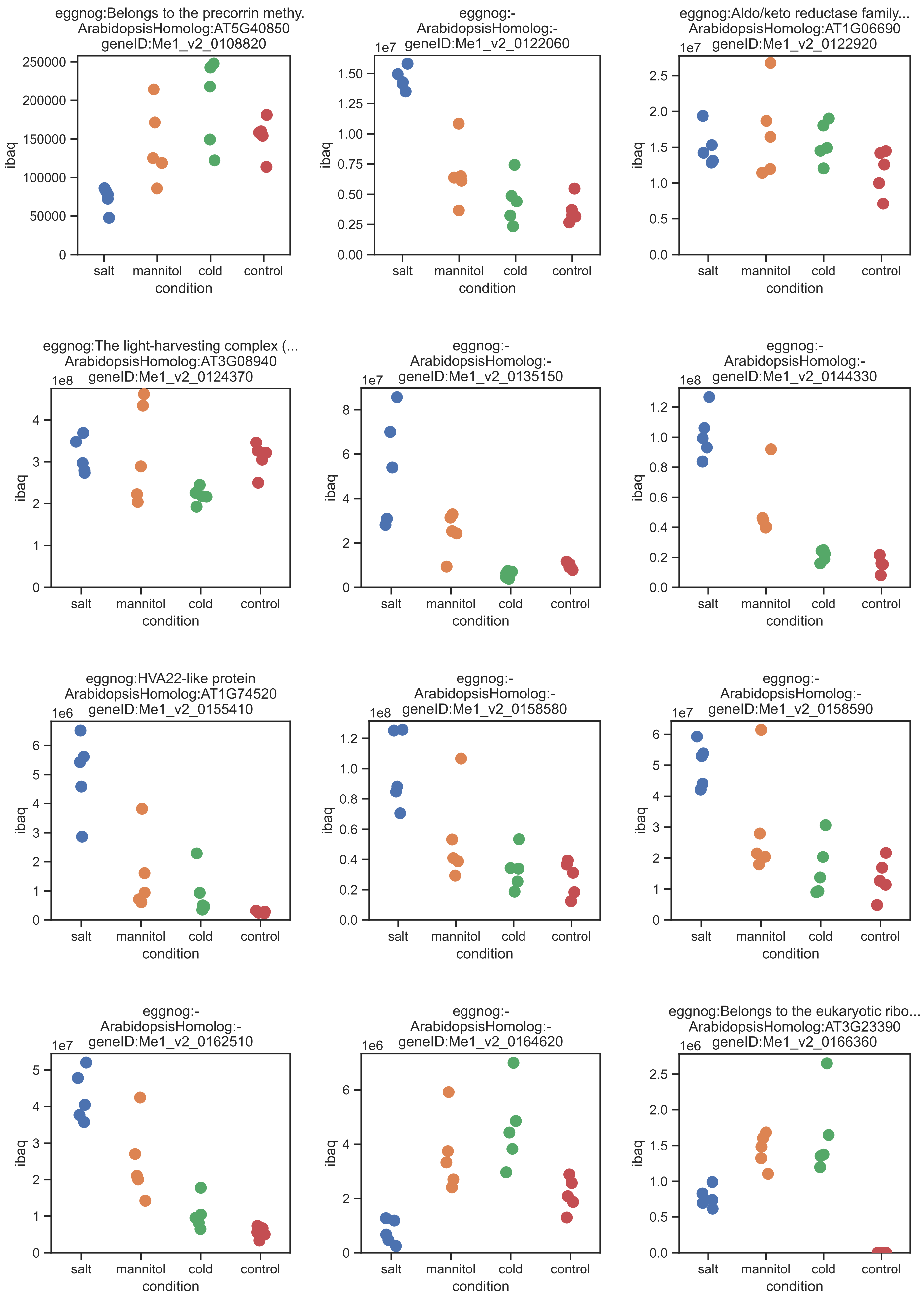
eggnog:Belongs to the precorrin methy.
ArabidopsisHomolog:AT5G40850
geneID:Me1_v2_0108820
250000
200000
150000
100000
50000
0
ibaq
salt
mannitol
cold
control
condition
eggnog:-
ArabidopsisHomolog:-
geneID:Me1_v2_0122060
1e7
1.50
1.25
1.00
0.75
0.50
0.25
0.00
ibaq
salt
mannitol
cold
control
condition
eggnog:Aldo/keto reductase family...
ArabidopsisHomolog:AT1G06690
geneID:Me1_v2_0122920
1e7
2.5
2.0
1.5
1.0
0.5
0.0
ibaq
salt
mannitol
cold
control
condition
eggnog:The light-harvesting complex (...
ArabidopsisHomolog:AT3G08940
geneID:Me1_v2_0124370
1e8
4
3
2
1
0
ibaq
salt
mannitol
cold
control
condition
eggnog:-
ArabidopsisHomolog:-
geneID:Me1_v2_0135150
1e7
8
6
4
2
0
ibaq
salt
mannitol
cold
control
condition
eggnog:-
ArabidopsisHomolog:-
geneID:Me1_v2_0144330
1e8
1.2
1.0
0.8
0.6
0.4
0.2
0.0
ibaq
salt
mannitol
cold
control
condition
eggnog:HVA22-like protein
ArabidopsisHomolog:AT1G74520
geneID:Me1_v2_0155410
1e6
6
5
4
3
2
1
0
ibaq
salt
mannitol
cold
control
condition
eggnog:-
ArabidopsisHomolog:-
geneID:Me1_v2_0158580
1e8
1.2
1.0
0.8
0.6
0.4
0.2
0.0
ibaq
salt
mannitol
cold
control
condition
eggnog:-
ArabidopsisHomolog:-
geneID:Me1_v2_0158590
1e7
6
5
4
3
2
1
0
ibaq
salt
mannitol
cold
control
condition
eggnog:-
ArabidopsisHomolog:-
geneID:Me1_v2_0162510
1e7
5
4
3
2
1
0
ibaq
salt
mannitol
cold
control
condition
eggnog:-
ArabidopsisHomolog:-
geneID:Me1_v2_0164620
1e6
6
4
2
0
ibaq
salt
mannitol
cold
control
condition
eggnog:Belongs to the eukaryotic ribo...
ArabidopsisHomolog:AT3G23390
geneID:Me1_v2_0166360
1e6
2.5
2.0
1.5
1.0
0.5
0.0
ibaq
salt
mannitol
cold
control
condition

**Supplementary Figure** 4.
Total Proteome

**A.** Fraction of the total SnRK2/PYL/PP2CA/ABF protein abundance in *Mesotaenium endlichrianum* based on the iBAQ values.

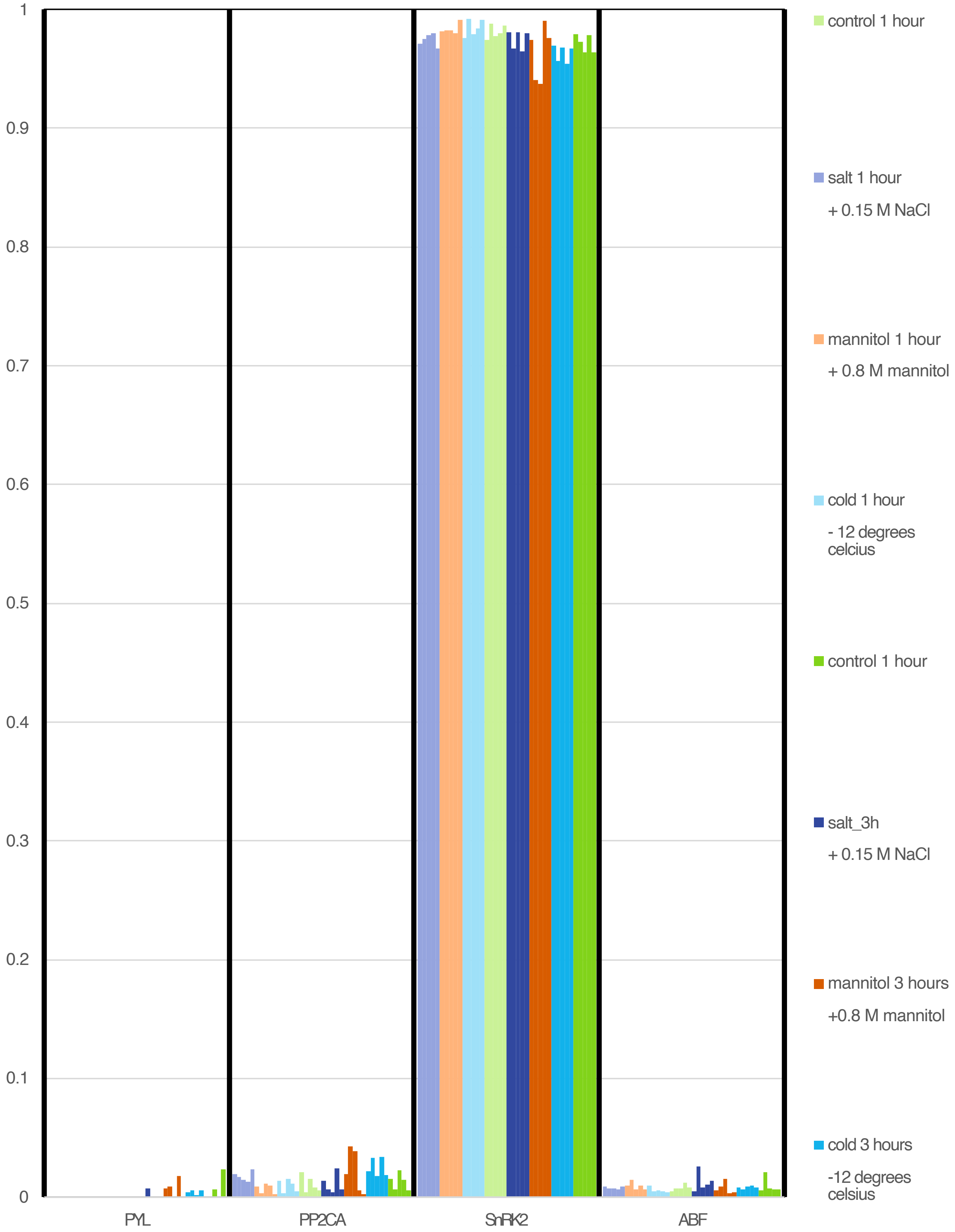
